# Linking H3K36me3 to transcriptional dynamics

**DOI:** 10.64898/2026.10.12.755748

**Authors:** Lawrence Hawkins, Shuo Wang, Jing Yuan, Massimo Cavallaro, Daniel Hebenstreit

## Abstract

Genes are transcribed in bursts as they fluctuate between on and off states, in a process modulated by histone modifications. The mammalian histone methyltransferase SETD2 is solely responsible for depositing H3K36 trimethylation, which recruits histone deacetylases and regulates histone occupancy. Nucleosomes are known to act as a barrier to transcription elongation and have been observed to impact elongation rate. H3K36me3 has been associated with long pausing sites in gene bodies and is implicated in reducing speed of successive RNA polymerase II (Pol II). However, the role of SETD2 and H3K36me3 in transcriptional dynamics are unknown. Here we show upon SETD2 knockout and H3K36me3 depletion in murine C2C12 myoblast cells that overall gene expression is reduced and splicing fidelity affected, as 5’ mutually exclusive exons are favoured over 3’ ones. Furthermore, transcriptional burst size is reduced, while bursting frequency increased. A simple mathematical model suggests that the observed changes in transcriptional dynamics can be explained by less stringently controlled polymerase pausing in the gene bodies of H3K36me3 depleted cells.

## INTRODUCTION

Since the early observations of transcriptional bursting in the 70s, understanding its regulation has been a major area of interest [1]. It is now understood that the burst-like activation of genes from ‘off’ to ‘on’ is a tightly regulated process, whereby a burst can span from tens of minutes to over an hour in mammalian cells [2]. Regulation of gene expression is complex and can involve modulation of initiation, elongation, and termination stages. To rapidly respond to stress and developmental signals, chromatin remodelling proteins establish chromatin states to augment transcriptional programmes via transcriptional burst size and frequency to modulate overall gene output [3]. While recent work in single cell sequencing and imaging approaches have begun to reveal the complexity of transcriptional bursting, to untangle the complex signalling between epigenetic modifications and transcription machinery requires a comprehensive systems-level approach.

A critical challenge in transcriptional bursting research is the lack of a consensus model to describe the dynamic switching between transcriptionally active and inactive states. In current models, different levels of transcriptional activity, or states, are typically assigned to gene promoters to explain their pulsatile nature [4]. Single state models have been used where genes are in a permanent ‘on’ state, which gives rise to a Poisson distribution of RNA counts. Although these models have been shown to fit count distributions for some highly expressed yeast genes, they are unlikely to account for the complex regulation of many genes in higher eukaryotes [5]. The more popular two state model assumes that genes can exist in either ‘on’ or ‘off’ states. This model explains the dynamic output of many genes well and is widely used to study the systems that regulate transcription [6–9]. It assumes a constant transcription rate while the gene is in the ‘on’ state, with stochastic switching between the ‘on’ and ‘off’, thus resulting in bursts of mRNA output followed by periods of inactivity. This model mathematically encodes for two parameters that describe the bursting: the burst size (mRNA produced per burst) and the burst frequency. Besides providing insight into the single cell heterogeneity in transcription, the estimation of these two parameters allows us to investigate the role of regulatory features, like chromatin states and epigenetic modifications.

Techniques involving fluorescent reporters and RNA count data have been widely used to study transcriptional bursting. MS2 assays use protein reporters to bind engineered sequences in nascent RNA transcripts to reveal transcriptional dynamics in real-time [10]. Single cell RNA-seq (scRNAseq) generates a genome-wide snapshot of transcript counts across a population of cells. Similarly, the smFISH technique employs fluorescent probes to measure transcript abundance in fixed cell populations and also produces count data [11]. The cell-to-cell variability in gene expression data obtained from these experimental setups can then be used to estimate bursting parameters by fitting it to mathematical models. Particularly with scRNA-seq datasets, this has become a powerful strategy to probe regulatory circuits controlling the bursting of every gene.

Histone modifications form complex signalling networks to regulate chromatin activity and consequently gene expression. Histone subunits can be methylated or acetylated at several lysine residues [12]. Single lysine residues can be mono-, di-, tri-methylated, or acetylated. Methyl marks are added to histones by Histone Methyltransferases (HMTs) and removed by Histone demethylases (HDMTs). Acetyl marks are added by Histone Acetyltransferases (HATs) and removed by Histone Deacetylases (HDACs). Acetylation promotes active chromatin by weakening histone-DNA interactions, which facilitates transcription factor recruitment. Methylation can be activating or repressive, depending on the histone residue it is deposited on and its level of methylation. Distinct methylation modifications alter the histones’ conformation and consequently their affinity to transcription factors and chromatin remodelers. Control of regional histone modification profiles is essential to regulating chromatin architecture and transcriptional activity [3].

For instance, repressive states such as constitutive heterochromatin and facultative heterochromatin are defined by H3K9me3 and H3K27me3, respectively. Enrichment of these marks is associated with gene repression, whereas profiles of acetylation among other marks indicates gene expression and euchromatin regions [3]. Active enhancer states are often defined by H3K4me1 and H3K27ac enrichment, while H3K4me3 is associated with promoters and CpG islands [13, 14]. In fact, both acetylated H3 and di-and tri-methylated H3K4 correlate with DNaseI sensitive chromatin [15]. While all acetylation typically associates with euchromatin, H4K12ac associates with newly assembled histones and H3K18 hypoacetylation is known to silence tumour suppressor genes in cancer [16, 17]. Downstream of promoters, active elongation states are primarily enriched for H3K79me2/3 and H3K36me3. Notably H3K79me2/3 decrease in enrichment from the transcription start site (TSS) to the transcription end site (TES), whereas the opposite is true for H3K36me3 [18]. Each histone modification plays a role in regulating chromatin state function and transcription. Notably precise control of epigenetic profiles regulates the energy requirement to traverse nucleosome barriers deposited throughout the gene [19–23]. Interestingly, Set2 deposited H3K36me3 in yeast is involved in regulating histone turnover [23, 24].

SET domain-containing 2 (SETD2) is an HMT that methylates H3K36 co-transcriptionally in the 3’ region of gene bodies [25, 26]. In mammalian cells, several other HMTs are responsible for H3K36 methylation: nuclear receptor-binding SET domain (NSD) protein 3 catalyses only H3K36me1; NSD1, NSD2, ASH1L and MYND domain-containing 2 (SMYD2) each catalyse H3K36me2; SETD2 alone catalyses H3K36me3 and is also capable of depositing H3K36me1/2 [27]. Notably, removing SETD2’s catalytic SET domain has been shown to impair H3K36me3 but not H3K36me1/2 deposition [28, 29]. Set2 is conserved between yeast and mammalian cells, however there are functional discrepancies. It is generally accepted that, as in yeast, mammalian SETD2 is co-transcriptionally recruited to elongating RNA polymerase II at the 3’ end of gene bodies. However, recent work has shown nuances to mammalian recruitment of SETD2; observations show SETD2 may recruit to Pol II during splicing, resulting in enrichment of H3K36me3 around exons and intron-containing genes [30]. In yeast, this is co-ordinated by a battery of enzymes coordinating a shift from serine 2 to serine 5 phosphorylation downstream of the transcription start site (TSS) which increases Pol II affinity for Set2 (as reviewed in [31] and [32]). To highlight this distinction, [30] also shows that, in mammalian cells, H3K36me3 enrichment aligns better with the second exon downstream of TSSs rather than enrichment for serine 5 phosphorylation.

Pol II elongation dynamics and histone modifications are related also through the phenomenon of polymerase pausing, the transient stoppage of Pol II within genes. It occurs most prominently proximal to promoters, where it is linked to H3K4me3 modifications [33], but is also observed at other intragenic regions, where it occurs for various reasons, including the barriers formed by nucleosomes and/or histone marks on these, splicing, and others [34–36].

SETD2 is arguably the only HMT responsible for depositing H3K36me3 in mammalian cells and is known to have roles in regulating acetylation, histone turnover, splicing, and DNA repair. H3K36me3 recruits HDAC1/2 in mammalian cells via PWWP2A, which binds the modification [37]. Further, murine H3K36me3 has been shown to promote DNA methylation of active gene bodies [38]. Both deacetylation, which increases affinity between DNA and histones, and DNA methylation functions of H3K36me3 are believed to suppress spurious transcription initiation. Further, H3K36me3 recruits Isw1b in yeast to reassemble histones in the wake of the transcription fork protecting histone modifications in the gene body [23]. Together with HDACs and Isw1b, H3K36me3 appears then to strengthen the DNA-histone interaction and reassemble disrupted histones, protecting histone modifications during transcription. In fact, loss of Set2 in yeast has been shown to increase histone turnover [24]. We previously observed long Pol II pausing sites overlapping with H3K36me3 enrichment and hypothesised that H3K36me3-dependent hypoacetylation promoted ‘speed bump’-like regions dense in nucleosomes [39, 40]. Interestingly, recent studies in mammalian cells have linked H3K36me3 and the co-transcriptional deposition of m^6^A modification to mRNA via METTL14 [41]. This is notable, since more slowly transcribed genes have been seen to exhibit less promoter proximal-pausing, greater H3K27me3/H3K4me3 fraction at promoters, and m^6^A deposition on nascent mRNA [42]. That H3K36me3 may have a key role in m^6^A deposition and that this mark is enriched at slower transcribed genes, indicates a potential relationship between the histone mark and elongation rate. Other roles for H3K36me3 include suppressing facultative heterochromatin associated H3K27me3 [43]. Although Set2 in yeast is known to only bind histones, ongoing research has found mammalian SETD2 is more promiscuous [44]. Recent work has shown SETD2 methylates the cytoskeletal protein α-tubulin and transcription factor STAT1 [45, 46]. These findings highlight a challenge in discriminating SETD2 loss from H3K36me3 depletion.

The precise role of histone modification in transcriptional bursting is unclear, however H3K36me3 has been linked to transcriptional pausing and regional hypoacetylation [3, 40, 47–54]. We seek to understand how SETD2 loss and consequent H3K36me3 depletion of chromatin states affect transcriptional bursting. To address this we use a previously published SETD2 KO in C2C12 murine myoblast cells and combined analysis of scRNA-seq and, ChIP-seq data, along with modelling approaches to dissect these effects [55]. Our study revealed an overall downregulation of RNA expression in KO cells and a modulation of bursting dynamics in genes associated with H3K36me3. An analysis of splicing and a mathematical model suggests the altered bursting to be due to increased dispersion of pausing times in gene bodies.

## RESULTS

### Characterising SETD2-/-Knockout C2C12 murine cells

Our previous findings indicated that, although H3K36me3 marks are associated with elongation, we found these to correlate with long polymerase pausing times in gene bodies [40]. Our goal was thus to study the effects of H3K36me3 on the dynamics of transcription, in particular bursting behaviour. To this end, we used a previously published double SETD2 knockout C2C12 cell line that has strongly reduced H3K36me3 deposition [55]. A western blot against H3K36me3 confirms the knockdown efficiency (Fig. 1A).

**Figure 1.**
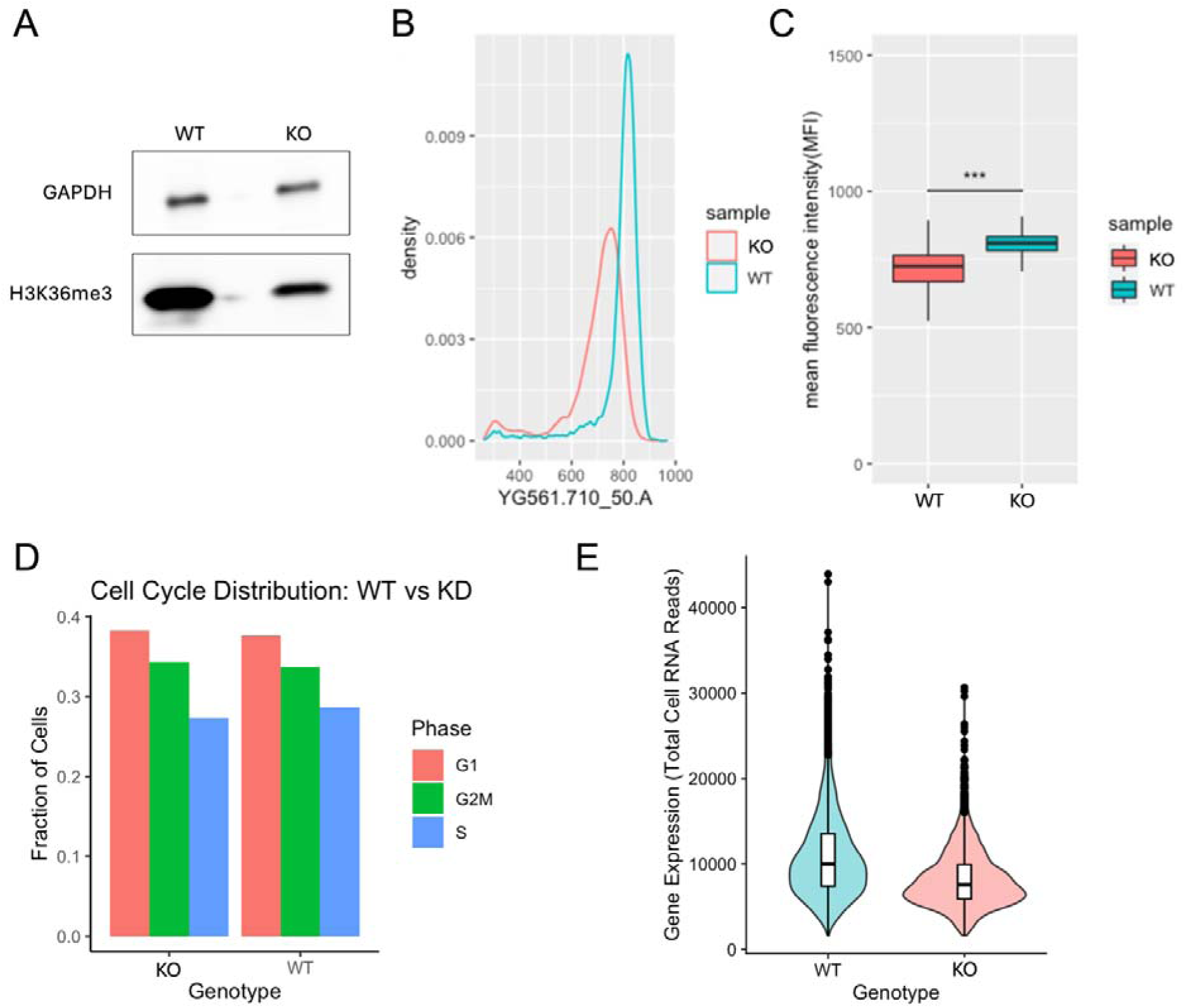
Characterization of global SETD2 knockout effects in C2C12 cells. **A**, Western blot detection of H3K36me3 and GAPDH as loading control in wildtype (WT) and knockout/down (KO) cells. **B**, distribution of total RNA content in single WT and KO cells as determined by flow cytometric analysis of fluorescent intensity corresponding to Pyronin Y. **C**, Distribution of mean of repeats of B. ***, p < 0.001, unpaired t-test. **D**, estimated fractions of KO and WT cells in cell cycle phases based on scRNA-seq data. **E**, MA Violin plot showing gene expression in KO vs WT cells.

As a first test of transcription in absence or presence of SETD2, we carried out a pyronin Y stain for total RNA. We evaluated results with flow cytometry, which demonstrated significantly reduced RNA expression in knockout (KO) compared to wild-type (WT) cells (Fig. 1B, C).

We then proceeded to carry out scRNA-seq on WT and KO cells as this allowed us to analyse gene expression in genome-wide fashion and analyse transcriptional dynamics based on single-cell RNA distributions. Based on genes known to be expressed in patterns related to the cell cycle, we estimated the proportion of cells in different cell cycle phases (Fig. 1D). This showed no obvious cell cycle defects in the KO cells, confirming our visual observations when culturing the cells. Inspecting global mRNA expression, however, reveals an 8.02% downregulation in overall expression levels in KO compared to WT, which agrees with our findings relating to total RNA (Fig. 1E). An MA plot is shown in Fig. S1.

### Investigating splicing in KO cells

The distribution of H3K36me3 marks reflects intron/exon structure [40] and the marks are implicated in splicing [40, 56, 57], which in turn relates to the dynamics of transcription [56, 58]. Plotting average H3K36me3 density in gene bodies based on external data for the cell type we were using (C2C12) confirms known profiles; the density increases towards the 3’ end of genes (Fig. S2A), is higher within the CDS (Fig. S2B-C), and depends on intron/exon boundaries, as reflected by higher signals in successive exons and introns (Fig. 2). We note that this increase is not smooth and appears somewhat step-like, in particular at the second and third exons and introns, as confirmed by two different datasets (Fig. 2). Displaying the same data as heatmaps reveals a striking pattern for exonic H3K36me3 marks, whose transcriptome-wide densities transition from over-to under-representation compared to adjacent regions (Fig. 3). While differential enrichment of H3K36me3 on expressed exons has been described [56], a depletion appears to be a new finding. Repeating this analysis for genes selected for different expression levels conserves these patterns, suggesting that they are not due to absolute expression levels (Fig. S3A-B). We neither find evidence that the depletion is due to exclusion of exons; there is no positive correlation between H3K36me3 signal and RNA read densities when comparing upstream and downstream mutually exclusive exons (MXE; Fig. 3B). Note that we considered only MXE with more RNA in the upstream exon as a conservative means to avoid capturing a potential bias stemming from the poly-A-priming of the RNA-seq library.

**Figure 2.**
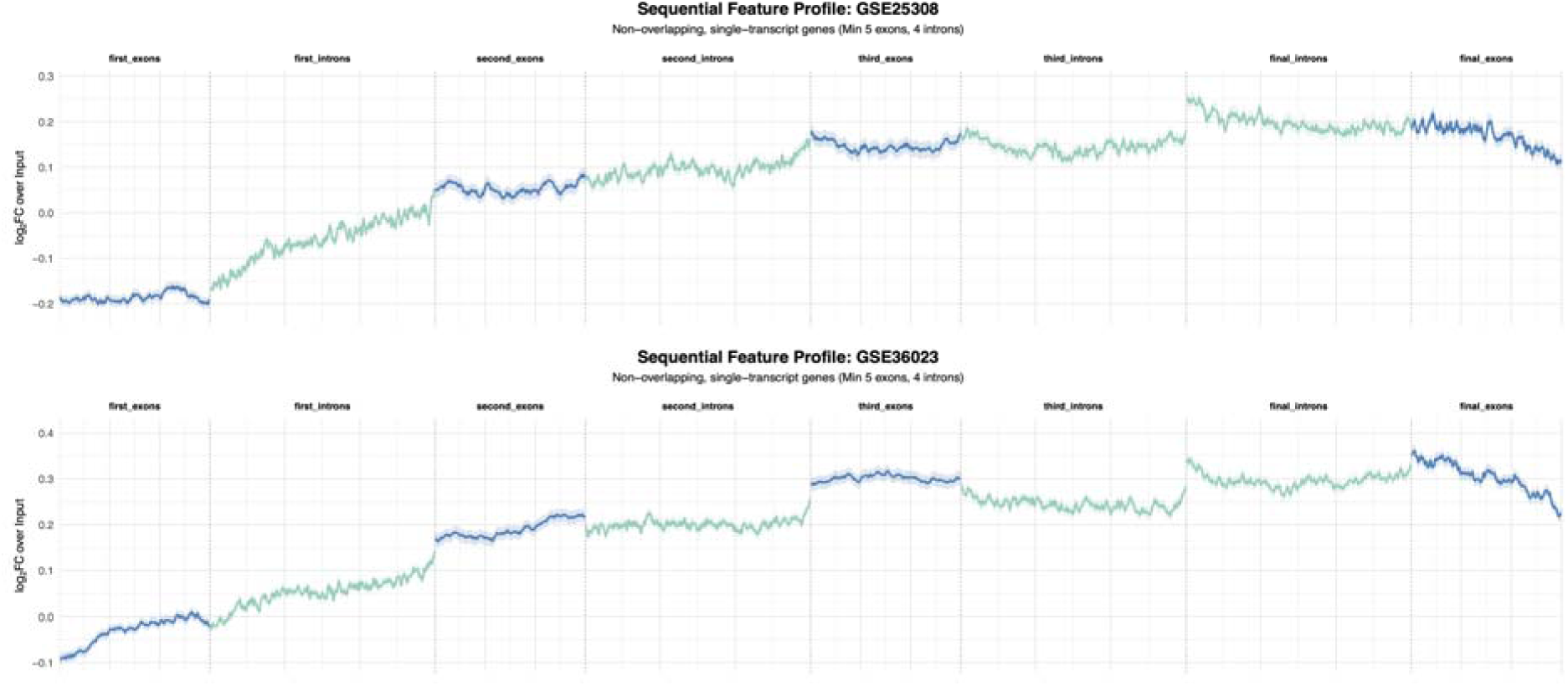
Average intragenic H3K36me3 density. Average densities of H3K36me3 marks in 1^st^ to 4^th^ exon (blue) and intron (green) relative to inputs for two different datasets as indicated.

**Figure 3.**
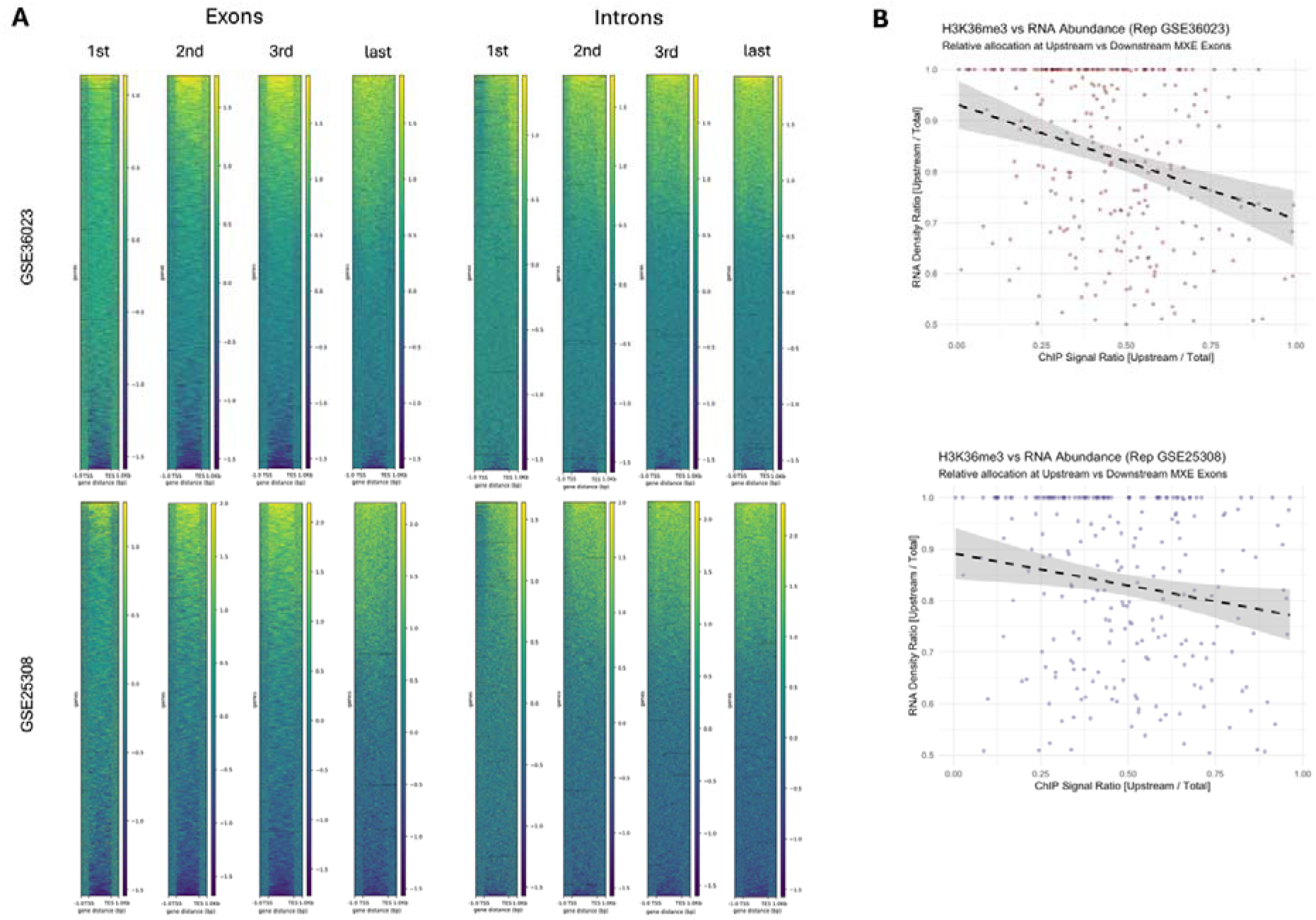
Intragenic H3K36me3 distributions. A, **d**ata as in figure 2, shown as heatmaps that resolve individual transcripts. **B**, mutually exclusive exons were identified and the signals at the upstream and downstream exon was calculated for H3K36me3 marks and RNA-seq read densities. Pairs with lower upstream than downstream RNA-seq read densities were filtered out to avoid capturing a potential bias of the poly-A-primed RNA-seq library. Ratios of upstream and total signal for each pair are plotted. r =-0.289, P = 3.298e-06 and r =-0.196, P = 0.002773 for top and bottom plots, respectively.

We next quantified the ratio of intronic and exonic reads for our scRNA-seq data in WT and KO cells to gauge the global splicing efficiency. This resulted in a significant but small difference (Fig. 4A; p=0.0007, Mann Whitney U test). However, a bigger effect of the knockout emerges when resolving splicing events in greater detail; 5’ MXE are >4 times more common in KO than WT cells (Fig. 4B, C). This suggests a link to Pol II’s elongation dynamics, as the splice machinery has longer to assemble on the 5’ MXE compared to the 3’ due to the direction of transcription [56, 58]. This may lead to favouring inclusion of the 5’ exon in KO cells, whereas in WT, H3K36me3 in exon 2 might stall Pol II, allowing time for the spliceosome to form on exon 2 to outcompete exon 1.

**Figure 4.**
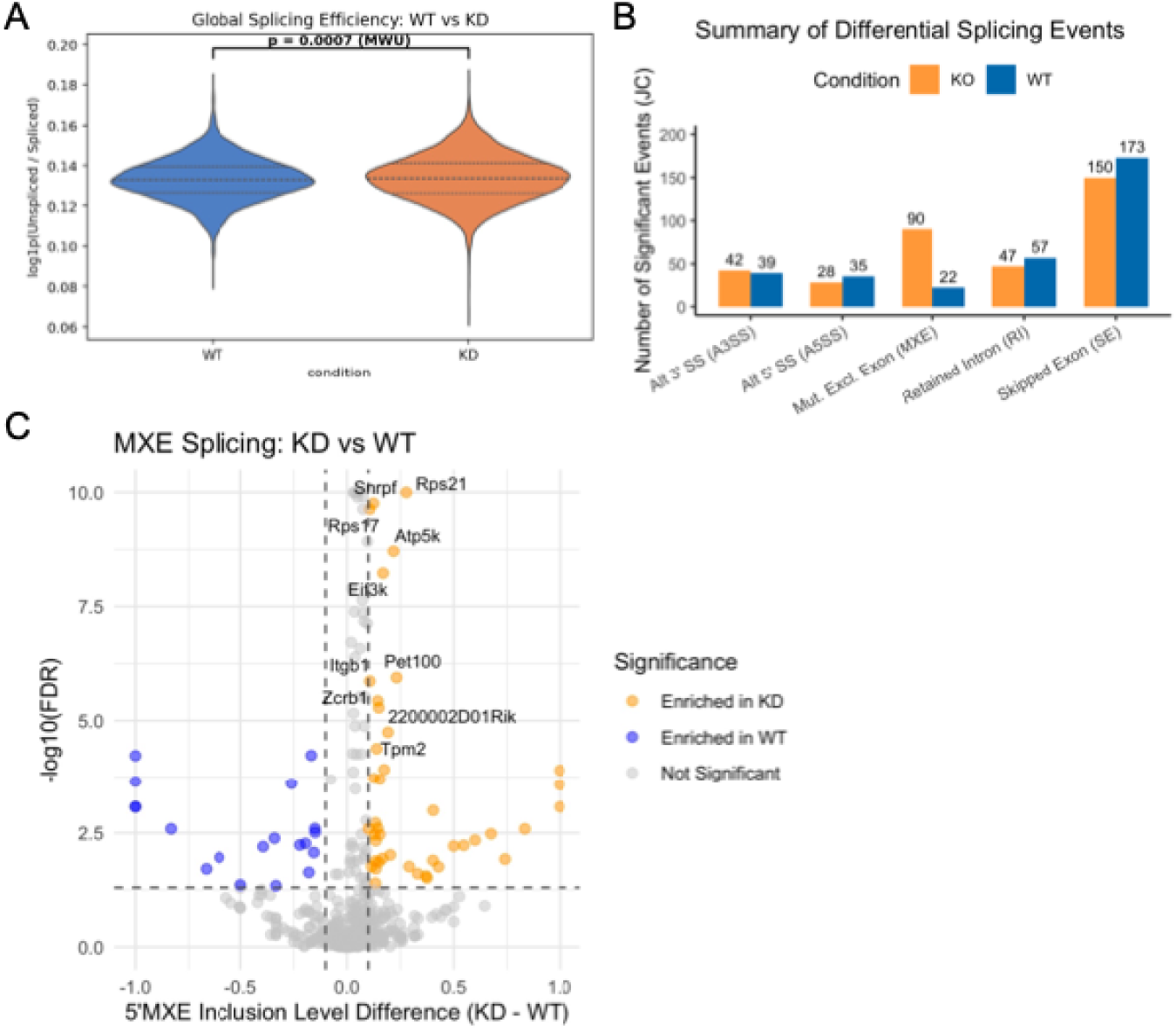
Splicing defects in SETD2 knockout cells. **A**, global splicing efficiency as measured by ratios of intronic and exonic reads in transcripts from WT and SETD2 KO cells. KO > WT, p=0.0007, Mann-Whitney test. **B**, Quantitation of types of splicing events in WT and SETD2 KO cells, showing increased utilization of 5’ compared to 3’ mutually exclusive exons in the latter. **C**, Volcano plot for inclusion level difference of 5’ mutually exclusive exons in WT vs SETD2 KO cells. All results based on scRNA-seq data.

### Characterizing chromatin states

Building on these results, we sought to augment our analysis with data for additional, common histone modifications. We identified 13 publicly available ChIP-seq datasets for histone acetylation and methylation in C2C12 myoblasts (Table S1) and confirmed agreement of the marks’ profiles with literature by plotting metagenes of their signal relative to input controls (Fig. S4).

To take account of redundancies and complementarities among these datasets, we used the ChromHMM tool [59] to learn chromatin states based on different combinations of these. To optimise the number of states we assessed the balance between capturing biologically important information and overfitting of the model (see Methods). We plotted Akaike’s Information Criterion and the Bayesian Information Criterion to understand at which number of states increasing the model size further provides no improved return on information (Figure S5A). In addition, to compare the redundancy of states we calculated the correlation between the states of a reference model and the states of every other model (Figure S5B). Interpretation of these metrics alongside the ChromHMM output supports the use of an 8-state model (Fig. 5A) [60].

**Figure 5.**
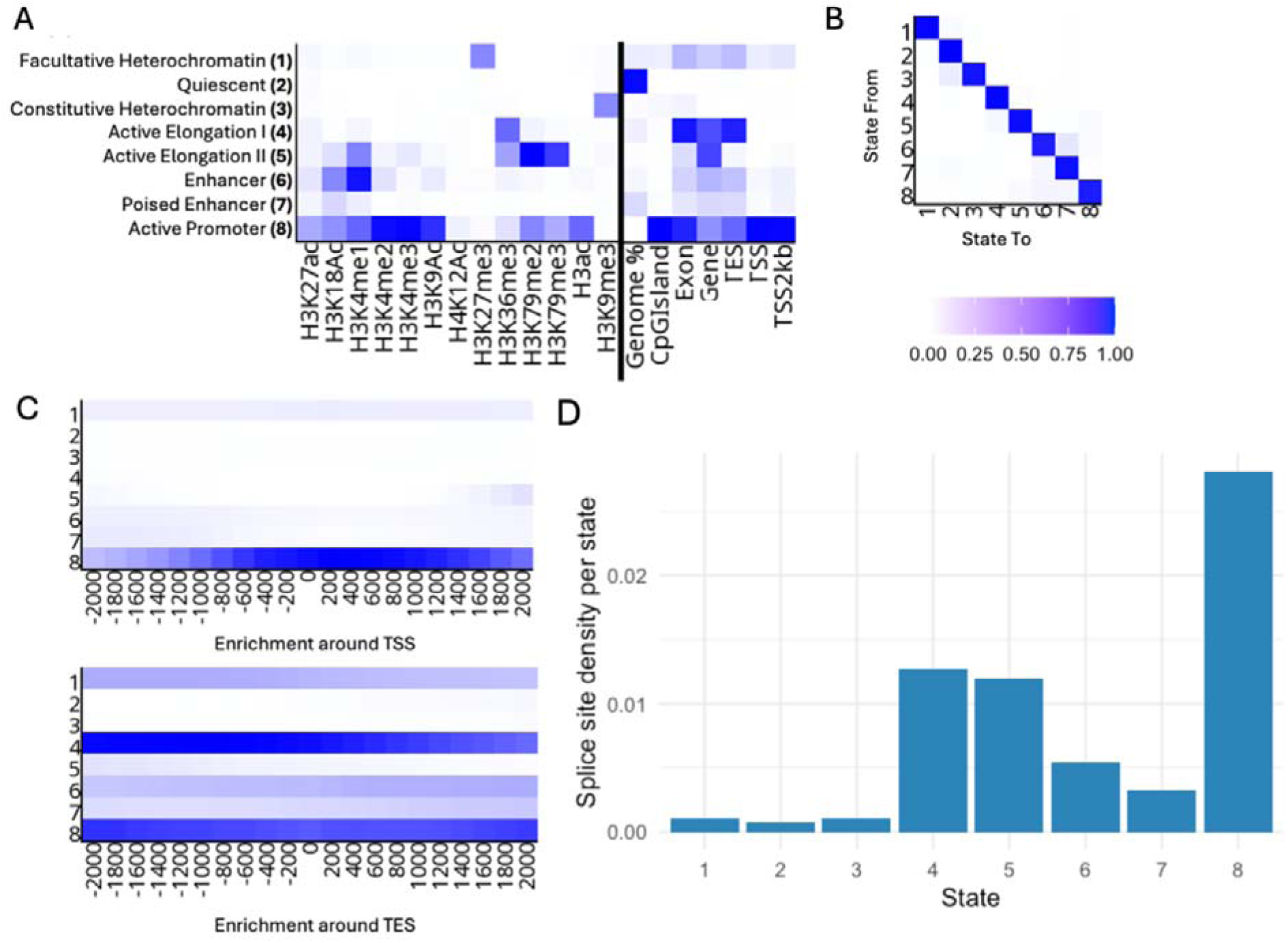
Chromatin state inference. **A**, allocation of 8 chromatin states with ChromHMM [REFXXX] based on datasets as indicated. State compositions and locations as indicated. **B**, state transition probabilities. **C**, state densities at TSS and TES. **D**, density of splice sites in the states. Blue colour in A to C indicates probability.

To interpret the functional role of each ChromHMM state we assessed their emission probabilities and regional enrichment scores (Fig. 5B, C). The profiles of states 1-3 denote transcriptionally silent or repressed chromatin states. State 1 shows a strong enrichment in H3K27me3 alone, indicating a repressed polycomb-regulated region, whereas state 3 is enriched for H3K9me3, denoting constitutive heterochromatin. State 2 describes regions enriched for none of the marks described in our model and functionally silent or quiescent chromatin. In contrast, states 4-8 describe profiles of histone modifications linked to active transcription. State 4 and 5 describe distinct states, both enriched for H3K36me3, H3K27ac, H3K4me1 and H3K79me2 all of which are linked to regions of active transcription [61]. Notably, state 4 shows weaker association to all these marks apart from H3K36me3 which is most strongly enriched here. State 5 is also enriched for H3K79me3 and H3K4me2-3, indicating a region of high transcription elongation. Interestingly, the distinct regional enrichments for state 4 and 5 modification profiles indicate independent regulatory roles. State 4 likely has an important regulatory role downstream in the gene body towards the TES as well as exons, whereas 5 is more involved in regulating elongation in intronic regions and immediately after the promoter (Fig. 5A, C). States 6 and 7 are enriched for H3K27ac, H3K18ac, and H3K4me1, all characteristic of regulatory enhancers. Greater enrichment for H3K4me1-2 in state 6 compared to 7 indicates more active enhancers compared to the poised enhancers described by state 7. The final state 8, highly enriched for active histone modifications including acetylation and mono-, di-, and tri-H3K4 and H3K79 methylation. Taken together with its sole enrichment at the TSS, state 8 likely describes active promoter regions.

We next aimed to understand how gene architecture associates with each chromatin state. To do this, we decided to identify genes with greater than 80% coverage for a given state. Having selected ‘high coverage genes’ for each chromatin state, we plotted the distribution of gene lengths and exon number, length, and density (exon length divided by gene length), all of which differ significantly by state (Fig. S6, p < 0.001, one-way ANOVA).

While the promoter-like state 8 shows highest exon density (Fig. S6A), probably because only short genes (Fig. S6B) with few introns are 80% covered by this state, the state with highest H3K36me3 enrichment, state 4, is also highest in terms of exon numbers per gene (Fig. S6C, p < 0.0001, TukeyHSD post-hoc). This agrees with the mark’s role in splicing.

To further investigate the implications of our chromatin states in splicing, we identified active splice sites with Regtools [62]. We filtered for splice sites supported by a minimum of 3 reads in our scRNA-seq dataset, split the genome into consecutive 200 bp bins, and counted overlaps between states and splice sites in these. We observed that all active transcriptional states, 4-8, were enriched for active splice sites (p < 0.0001, Phyper hypergeometric test). Notably, the promoter state (8) showed the greatest splice site enrichment, followed by states 4 and 5 at approximately one-fold reduction and 6 and 7 exhibiting half that level of enrichment again (Fig. 5D). As expected, heterochromatin and quiescent states (1-3) show poorest enrichment for active splice sites.

### Gene expression and chromatin states

We expanded our classification of genes regarding their chromatin state coverage and grouped them into increasing intervals of 20%, with a category of 0% corresponding to any of the other states except the one under consideration. We revisited our analysis of gene expression levels and used our scRNA-seq data to investigate the effect of SETD2 knockout and consequent H3K36me3 abolishment on the different classes of genes. Coverage of states 4, 5, and 8 positively correlated with expression levels, while the rest showed the opposite behaviour (Fig. 6A).

**Figure 6.**
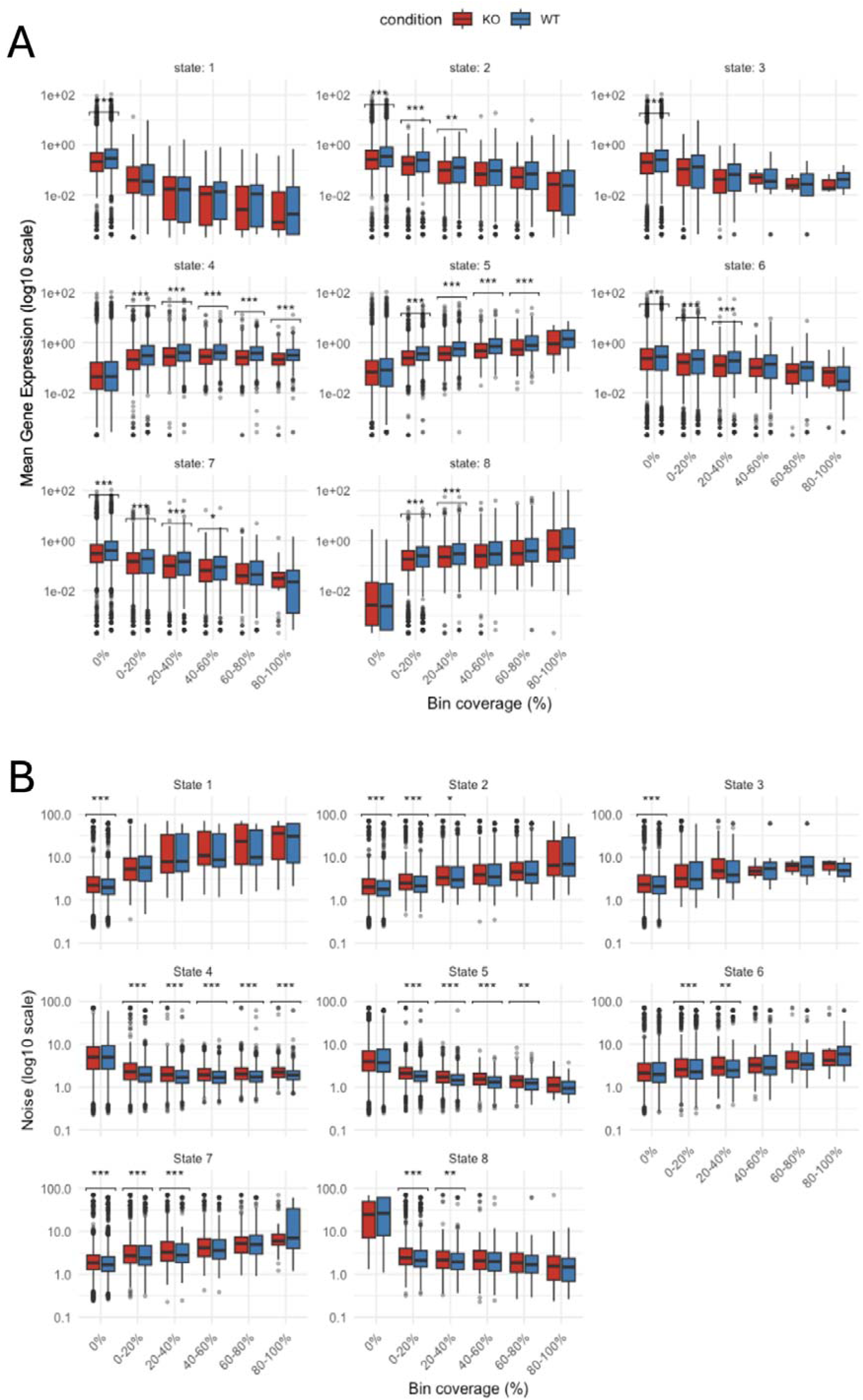
Effects of H3K36me3 loss on gene expression and noise, stratified by chromatin state coverage. Genes were classified according to their coverage by chromatin states 1 to 8 (Fig. 5). **A**, Mean expression levels were calculated based on UMI counts from scRNA-seq data. **B**, as A, for noise (CV^2^). Genotypes were compared using Mann Whitney U test with Bonferroni correction (* < 0.05, ** < 0.01, *** < 0.001). Numbers of genes sampled for WT and KO are n = 9167 and n = 8272, respectively.

The H3K36me3 enriched states, 4 and 5, exhibited the most notable decrease in gene expression (Fig. 6A). State 4 exhibited sole enrichment for H3K36me3 and showed that SETD2 loss caused a significant reduction in transcription at these genes (p < 0.001, Mann Whitney U test). Interestingly, genes with coverage greater than 0% and up to 80% for state 5 demonstrated significant reduction in transcription with the loss of SETD2 (p < 0.001, Mann Whitney U test. Although genes enriched in the heterochromatic states (1 and 3) showed no significant difference in expression in cells without SETD2, genes lacking these states showed a significant reduction (p < 0.001, Mann Whitney U test). The subset of genes with none of either heterochromatic state reveals a global downregulation of active genes upon H3K36me3 depletion. Enhancer-like states (6 and 7) and the promoter state (8) demonstrated similar trends; genes with low coverage greater than 0% (up to 40%) in these states showed significant reductions in gene expression (p < 0.001, Mann Whitney U test). We also plotted the transcriptional noise (quantified by the squared coefficient of variation squared, CV^2^), which shows similar, but inverted trends, confirming the frequently observed inverse correlation between mean and noise [63] (Fig. 6B).

We repeated these analyses with an alternative method of associating genes with chromatin states, this time stratifying genes by their 3’ most chromatin state. The results were similar, showing comparable overall trends and effects of the knockout (Fig. S7).

### Inferred transcriptional dynamics

Having observed subtle reduction in the gene expression levels following the loss of SETD2, particularly in states enriched with H3K36me3, we considered the effect of the mark on transcriptional bursting. Bursting dynamics were characterised by the overall mean expression, its CV^2^, the burst frequency and the burst size, all inferred from experimental scRNA-seq read distributions (M&M).

In WT cells, genes exhibited higher transcriptional bursting than in KO. An unadjusted comparison of the log of CV^2^ between WT and KO shows that WT genes tend to have slightly yet significantly lower noise than KO genes (Fig. 7A; p<2e-16, Mann Whitney U test), which is arguably due to higher overall expression levels in WT (Fig. 7A). In fact, by adjusting for the log of mean expression shows significantly higher noise in WT (lm, p<2e-16, t test). WT shows clearly larger log burst sizes than KO (Fig. 7A; p<2e-16, Mann Whitney U test), which holds when adjusting for differences in the log of mean expression (Fig. 7A; lm, p<2e-16, t test). This confirms that the larger burst size in WT is not simply due to higher average expression. Bursts also occur less frequently in WT than in KO (Fig. 7A; p<2e-16, Mann Whitney U test); consistently with the previous tests, adjusting for mean expression confirms that WT is associated with lower burst frequency (lm, p<2e-16, t test).

**Figure 7.**
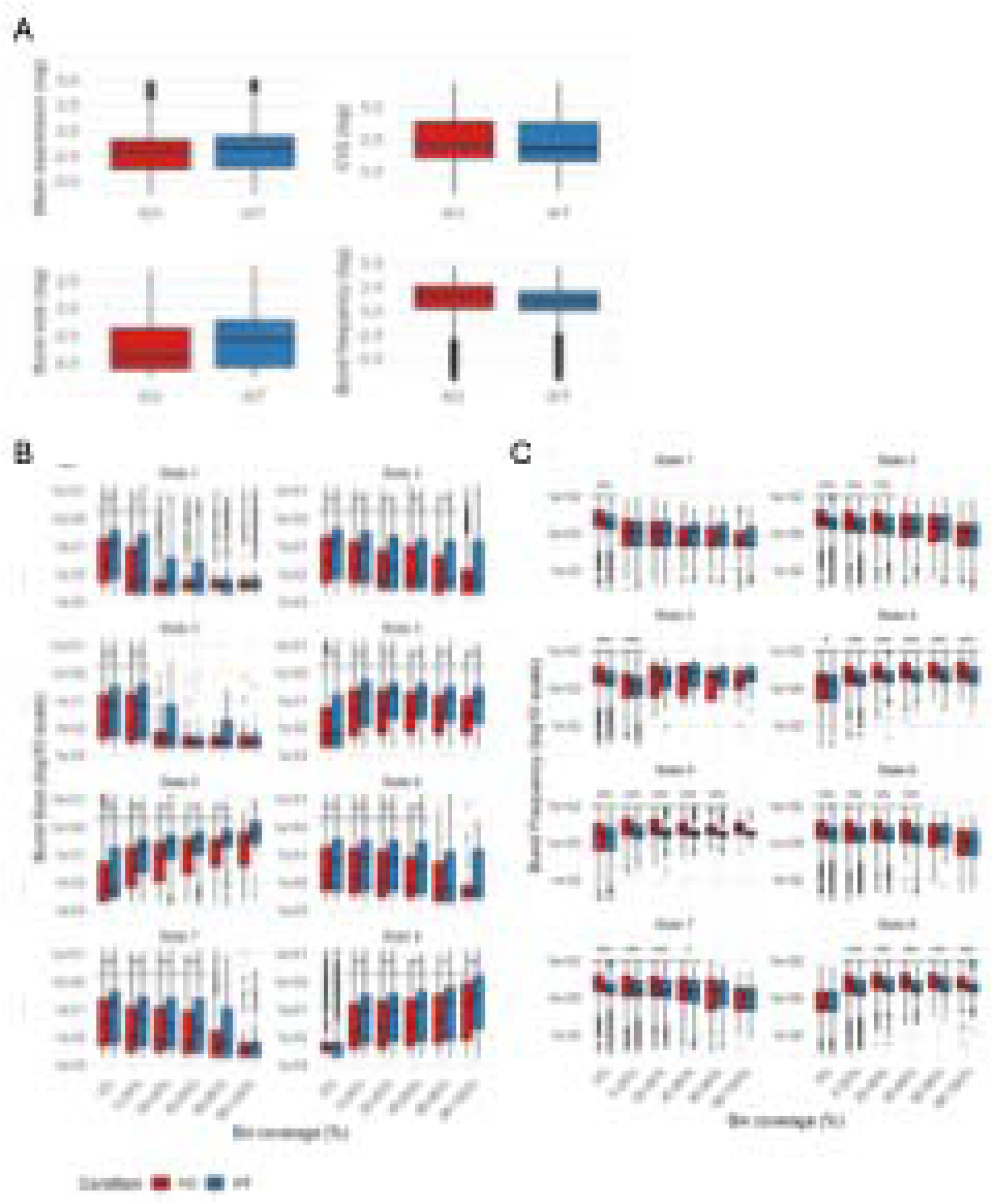
Effects of H3K36me3 loss on bursting parameters. A, mean expression, CV^2^, burst frequency, and burst size (= mean expression / burst frequency) were estimated from scRNA-seq count data using a Markov chain Monte Carlo (MCMC) approach. **B**, burst size, stratified by chromatin state. **C**, as B, for burst frequency. Genotypes were compared using Mann Whitney U test with Bonferroni correction (* < 0.05, ** < 0.01, *** < 0.001). For B and C, genes were classified according to their coverage by chromatin states 1 to 8 (Fig. 5).

We also assessed how chromatin state coverage modulates transcriptional bursting and how this relationship is affected by SETD2 loss. With depletion of H3K36me3, elongation states (4 and 5) show a significant reduction in burst size (Fig. 7B) and a mirrored increase in burst frequency (Fig. 7C) for all levels of coverage besides genes with 80-100% for state 5 (p < 0.001, Mann Whitney U test). It is worth noting that H3K36me3 is still important even at genes without these states (4 and 5), since they exhibited a significantly different distribution of burst size and burst frequency parameters (p < 0.05, Mann Whitney U test). Similarly, loss of H3K36me3 at genes with states 4 and 5 resulted in a significant reduction in expression and an increase in transcriptional noise (CV^2^) for all levels of enrichment besides 80-100% for state 5 (p < 0.01, Mann Whitney U test).

The enhancer like states (6 and 7) showed subtler changes across bursting parameters. Genes without enrichment of these states (6 and 7) and those with up to 60% coverage still showed significantly different distributions for each parameter after SETD2 loss (p < 0.05, Mann Whitney U test).

The promoter state (8) showed similar trends to the elongation states (4 and 5) and exhibited significantly reduced burst size and increased burst frequency compared to wild-type at all levels of coverage besides genes without this state (0% coverage bin) (p < 0.05, Mann Whitney U test). This is likely due to classifying genes without the promoter state effectively producing a group of transcriptionally silent genes, as demonstrated also by the mean expression values clustering at 0. Interestingly, only genes with no coverage or up to 40% coverage for the promoter state (8) showed a significant increase in CV^2^ (p < 0.05, Mann Whitney U test), indicating correlation between promoter state coverage and transcriptional noise. Alternatively, as expression and CV^2^ are generally inversely correlated, it may also reflect that higher promoter state coverage supports greater gene expression.

The repressive and quiescent states (1-3) show the least difference between WT and KO. Surprisingly, the quiescent state (2) shows a significant reduction in mean expression for all levels of coverage, including genes without this state (p < 0.01, Mann Whitney U test). However, expression does decline as coverage for the quiescent state (2) increases, likely due to genes becoming depleted in chromatin remodellers and transcription factors and thus transcriptional activity. The repressive states (1 and 3) show significant difference for each parameter only at low coverage, greater than 0% up to 20%, besides for burst frequency where the facultative heterochromatin state (1) shows no significant difference (p < 0.05, Mann Whitney U test).

Repeating these analyses for genes classified according to the 3’ most chromatin state largely recapitulates these findings (Fig. S8); the elongation states (4 and 5) and the enhancer like states (6 and 7) opposed each other in whether each bursting parameter was increased or reduced in respect to the promoter state (8). Both elongation states (4 and 5) show significant reductions in burst size and increases in burst frequency upon knockout (p < 0.01, Mann Whitney U test), while only state 4 shows a significant increase in CV^2^ and reduction in mean expression (p < 0.001, Mann Whitney U test). However, for burst size and burst frequency, all active states (4-8) and the quiescent state (2) show significant reductions and increases of similar magnitude, for both parameters (p < 0.01, Mann Whitney U test). Genes with the repressive states (1 and 3) as their final state show insignificant changes in bursting and expression with the loss of SETD2. Furthermore, genes ending in the elongation state 5 or the promoter state (8) show relatively smaller and not significant changes in mean expression or CV^2^ (Fig. S8).

### Model

We finally aimed to integrate our observations of altered bursting behaviour in elongation associated chromatin states into a model that can explain underlying mechanisms. We considered the total Pol II travel times from initiation to termination/mRNA release. These times are expected to influence the gene expression bursting patterns, with increased variability and dispersion leading to a flattening of the burst profiles, thereby reducing transcriptional noise. In particular, to account for reduced transcriptional noise in KO cells, we expect that the Pol II travel times in these cells (hereafter referred to as τ_ko_) are subject to more dispersion than the travel times in WT cells (τ_WT_), as illustrated in Fig 8A.

**Figure 8.**
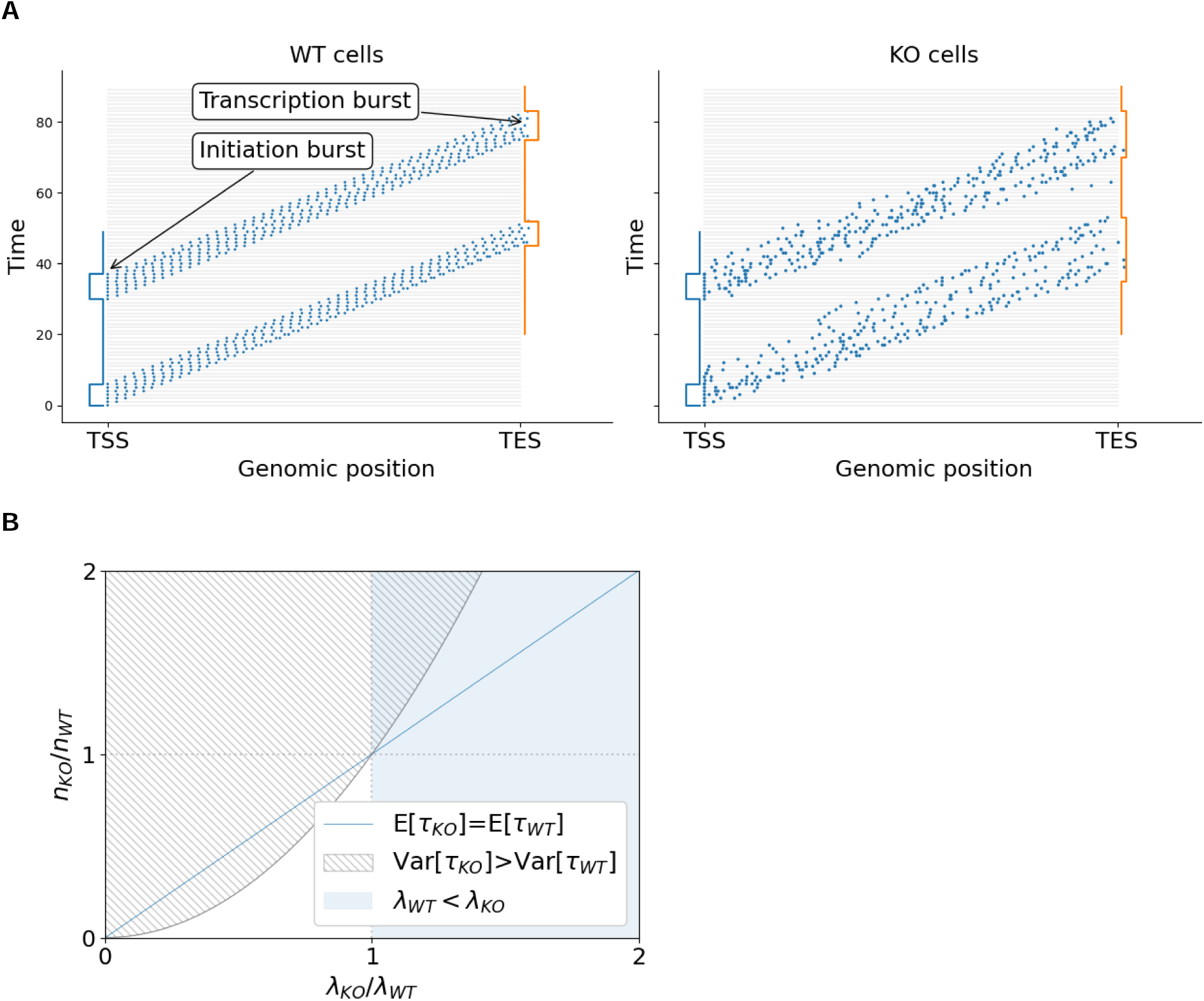
Model for Pol II dynamics. **A**, Schematic illustration of how trains of transcribing Pol II result in focused bursts in wildtype cells (left) and how reduced H3K36me3 levels add dispersion, reducing burst size and boosting frequency. **B**, Parameters are constrained to the intersection of shaded () and dashed () areas.

We assume that Pol II in a gene moves downstream through n successive discrete steps, each of random duration, independent of the other, and with an average duration 1/λ. According to this model, the average total travel time τ is simply given by E[τ] = n/λ, with variability quantified by the variance Var[τ] = n/λ^2^. In KO cell line genes, with reduced levels of H3K36me3, Pol II is subject to altered or diminished regulatory control, corresponding to a different number of discrete steps n_ko_ and inverse duration λ, compared with the number of steps n_WT_ and inverse duration λ in WT cells.

To explain the observed transcriptional bursting patterns, we require that Var[τ_ko_] > Var[τ_WT_], which implies 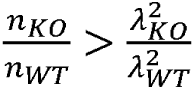 (dashed area in Fig 8B). Prior observations of prolonged pausing at H3K36me3-enriched sites [Zhang et al., 2021] also suggest that transcription in WT cells has longer effective step durations than in the cells where H3K36me3 is knocked down, encoded in the model as a smaller inverse duration (λ_WT_ < λ_ko_, shaded area in Fig 8B). Under these assumptions, it is worth noting that the allowed range (shaded and dashed area in the figure) is quite restricted and always falls above the horizontal line n_ko_/n_wt_ = 1 (thereby implying that n_ko_ > n_wt_) and above the diagonal line E[τ_WT_] = E[τ_ko_] (indicating faster elongation in WT). This suggests that, in WT cells, Pol II would progress through a smaller number of steps of longer duration, whilst keeping a lower total travel time from from initiation to mRNA release than in KO cells.

## DISCUSSION

Our results demonstrate that SETD2 and H3K36me3 have a role in regulating transcriptional bursting which impacts gene expression. We identify a chromatin state with strong H3K36me3 deposition, regionally enriched in exons and TESs.

Loss of SETD2 and H3K36me3 depletion resulted in significant reduction in overall gene expression. The reduction was strongest in genes covered by chromatin states with strong H3K36me3 enrichment. A similar observation in yeast noted that Set2 loss has a large impact on transcriptional networks [64]. Despite the overall reduction in expression some genes are overexpressed which supports a stabilising role for H3K36me3 in gene expression. Indeed, H3K36me3 has positive and negative effects on transcription; the mark antagonises PRC2 and facultative heterochromatin while suppressing cryptic transcription [37, 43]. The dual activity of H3K36me3 may be critical in establishing nucleosome barriers by regional hypoacetylation but antagonising further closing of the chromatin into heterochromatin with the deposition of H3K27me3. This activity may have a role in controlling the height of the nucleosome barrier Pol II needs to overcome, sufficiently high to stall Pol II but also low enough for transcription to traverse [19, 37, 40, 65, 66]. Depletion of H3K36me3 may thus dysregulate elongation through nucleosomes and significantly impair global transcription. We note that our results suggest that this reduction likely includes rDNA transcription, given that rRNA constitutes most cellular RNA and we find significantly lower amounts of total RNA in KO cells. This could be an indirect effect of rRNA transcription machinery downregulation or a previously unrecognized role for H3K36me3 and/or SETD2 in rRNA transcription. Further research will be necessary to explore this.

Chromatin remodellers other than SETD2 have been implicated in large scale transcriptome changes, too. Interestingly, H3K36me3 has been shown to coenrich with HDAC2 [37] and dual, but not individual inactivation of HDAC1/2, also shows large transcriptomic shift [67]. By depleting H3K36me3 and loss of HDAC2 targeting to genes, shifts in gene acetylation could lead to large shifts in histone turnover and gene expression. Indeed, both histone turnover and histone acetylation are frequently associated with gene activity [48, 68]. It would be interesting to see if SETD2 loss and H3K36me3 depletion is sufficient to dysregulate HDAC2 function and cause similar dysregulation of transcriptional programmes. It is notable however, that more genes were associated with upregulation than downregulation after HDAC1/2 KO, indicating a more complex picture [67]. Chd1 is a chromatin remodeller involved in nucleosome turnover, reassembly, and positioning [69]. In yeast, it is also involved in H3K4me3 and H3K36me3 organisation throughout the genome and loss of Chd1 leads to significantly reduced intron retention, highlighting its role in splicing [69, 70]. This factor is also associated with global reduction in transcription, which may indicate the importance of H3K36me3 and nucleosome organisation in maintaining gene expression[71]. Furthermore, the BAF or mSWI/SNF complex is also a chromatin remodeller which has roles in nucleosome positioning and opposing PRC2 and facultative heterochromatin spread [72]. Its subunits are frequently mutated in cancers and its loss of function leads to wide-spread changes in transcriptional programmes [73, 74]. Notably, in mice BAF binds hepatoma-derived growth factor-related protein 2 (HRP2), which in turn binds H3K36me2 [75]. Upregulation of BAF subunit DPF3a was observed during myogenesis and drive muscle differentiation. This highlights the importance of H3K36 methylation in the regulation of developmental transcriptional programmes, although it is worth noting that SETD2 loss is primarily responsible for H3K36me3 deposition alone [76]. Similar to BAF’s role in suppressing PRC2, H3K36me3 on nucleosomes has been shown to antagonise PRC2 activity [43]. This indicates the intricate signalling between chromatin remodellers especially in organising chromatin accessibility and heterochromatin borders. The closely overlapping roles for SETD2, BAF, Chd1, and HDAC1/2 in regulating nucleosome occupancy highlights the key role of these factors in controlling global gene expression.

We evaluated whether the impact of SETD2 loss was greater for genes with different levels of coverage for specific chromatin states, elevated Pol II density within those states, or depending on the chromatin state present at their TES. First, we assessed expression based on the coverage of genes for any chromatin state. We found that only genes with greater levels of coverage for H3K36me3 enriched states experienced a significant shift in gene expression after SETD2 loss, whereas genes covered by states without H3K36me3 enrichment were only significantly reduced after SETD2 loss if coverage was below 60%. This may be due to genes with greater coverage for chromatin states without H3K36me3 passing a threshold whereby the space left for H3K36me3 enrichment is insufficient to produce a significant effect on gene expression following depletion. We further note that the histone modifications we examined constitute only a subset of the known ones, but there is substantial redundancy {Swygert, 2014 #12}, even within our limited set, as 8 states appear enough to capture the features of 13 different types of marks.

We then grouped genes based on the chromatin state at their TES. The aim was to assess whether 3’ chromatin states have a role in altering gene expression. Previously, at the end of elongation rate-limiting steps 1 minute long have been observed along with increased Pol II density highlighting termination as an understudied topic in transcription [77, 78]. We observed similar trends in downregulation for all chromatin states associated with active transcription, though only significant reductions were observed for the chromatin state with strong H3K36me3 and the enhancer-like states with H3K4me1 and acetylation.

A limitation of classifying chromatin states by position is that they each have positional biases, showing differential enrichment around different genomic features. As a result, sample sizes for each category were uneven, giving different groups greater statistical power. For example, the chromatin state with strong H3K36me3 showed significantly stronger enrichment at TESs compared to all other states, which is also reflected by the fact that, when categorising genes by their final state, this category had almost 2-fold greater gene numbers than any other group.

We observed a SETD2-dependent reduction in burst size and increase in burst frequency which is accompanied by significant reductions in gene expression (Figure 7). Previously, we observed H3K36me3 enrichment to overlap with long pausing sites in the gene body and postulated that regional hypoacetylation and increased histone occupancy could have a ‘speed bump’-like effect on Pol II elongation [39, 40]. The role of H3K36me3 in reducing histone turnover and recruiting HDACs provide evidence for its role in establishing nucleosome dense barriers to slow elongating Pol II [24, 79, 80]. Our observation that H3K36me3 depletion reduces burst size of genes enriched for this mark indicates that stability of Pol II during elongation may be compromised. This may be a result of loss of chromatin remodellers to facilitate traversal of Pol II through nucleosome rich regions [23]. Here, we saw SETD2 loss affects burst size and burst frequency, while another study has observed the chromatin context of HIV promoter integration sites to affect only burst size [81]. This difference may be due to varying chromatin accessibility at promoter integration sites, while in contrast, co-transcriptionally deposited H3K36me3 likely produces a consistent modification pattern regardless of chromatin accessibility. This may also explain why other studies have observed chromatin context to not impact bursting [82, 83]. Perhaps H3K36me3 is critical to facilitating transcription through nucleosome dense chromatin, however upon depletion of this mark and recruitment of chromatin remodellers, bursting is significantly impacted.

Interestingly, models have shown that bursting can occur due to long pausing in the gene body, irrespective on initiation frequency [84]. Since we previously observed long pausing at H3K36me3 sites [Zhang Jie et al 2021] and burst sizes are mostly reduced upon SETD2 knockout, shorter Pol II pausing times are to be expected for the latter. This appears to agree with our splicing-related findings, where we find the 5’ MXE usage to be increased in KO cells, which is indicative of reduced time to select the alternative exon. Our mathematical model postulates a reduced coordination and/or less stringent regulation of pausing times or stalling of Pol II, which can explain the reciprocal shift in burst sizes and frequencies we observe. The model is agnostic to the underlying mechanisms but appears plausible in light of the hypotheses expressed above.

Here, we removed overlapping genes from our analyses, however work showing transcription of intragenic enhancers by Pol II may antagonise transcription of their host gene. This may implicate gene overlap as a potential transcriptional control [85, 86]. Previously, we hypothesised H3K36me3-dependent hypoacetylation stalls Pol II to potentially protect collision of successive Pol II [40]. Investigating how transcription at overlapping genes may affect transcriptional dynamics presents an exciting opportunity to study whether the epigenome may have a role in regulating Pol II’s elongation rate to protect against collisions [19, 21, 66, 87, 88].

More work is needed to understand the epigenetic changes that take place following H3K36me3 depletion and SETD2 knockout and whether they lead to major chromatin state reorganisation. We anticipate that SETD2 knockout has negligible effects on bulk mono-and di-methylated H3K36 deposition, as shown previously [28, 29]. However, revealing organisation of chromatin states in absence of H3K36me3 will help understand the role of this mark in organising chromatin states and the effect of its loss on transcription. Indeed, if there is a bias for one histone modification to replace H3K36me3, it may help explain the effect of SETD2 loss on transcription. Further work will be needed to directly measure the rate of nascent mRNA transcription after SETD2 knockout. Understanding the relationship between elongation rate and levels of gene body H3K36me3, acetylation and histone occupancy should be the focus of future investigations.

In conclusion, we demonstrate that SETD2 and H3K36me3 have roles in amplifying burst size and reducing burst frequency which significantly impacts overall gene expression. This supports a model for H3K36me3 in maintaining a nucleosome dense speed bump, slowing successive Pol II [40]. By stabilising elongating Pol II, we expect burst size is maintained reducing the burst frequency required to reach appropriate mRNA output. We speculate that, without H3K36me3 enrichment downstream in the gene body, long-pausing sites are abolished and Pol II stall more frequently, potentially leading to slower elongation together with more diffuse, smaller, and frequent bursts, as captured by a mathematical model.

## MATERIALS AND METHODS

### Establishing efficacy of SETD2-/-knockout and H3K36me3 depletion

Flow cytometry was carried out to assess population and RNA content of wild-type (WT) murine C2C12 cells compared to a previously published SETD2-/-knockout C2C12 cell line [55, 89]. 1×10^6^ WT and KO cells were harvested at 80% confluence, undergoing a PBS wash prior to staining at 1×10^6^ cells/ml. Fixed cells were treated with 5 μl of Pyronin Y (100 μg/ml, Sigma) then incubated in the dark at 37°C for 15 minutes. Prior to incubation, cells were on ice and covered during the experiment. Flow cytometer (BD Fortessa) measurements were taken at approximately 800 events/second. A dot plot was used to assess cell dimensions and exclude doublets and clumps.

To assess efficacy of H3K36me3 depletion, C2C12 cells (WT and KO) lysed in RIPA buffer and protease and phosphatase inhibitor (Sigma). Protein concentration was determined using PierceTM BCA Protein Assay Kit (ThermoFisher). 15 mg of protein was allowed to run on precast 4% SDS-PAGE gel (abcam) and transferred to nitrocellulose membrane. The membrane was incubated with H3K36me3 (abcam134391) and GAPDH (Proteintech) antibodies followed by secondary antibodies, one anti-rabbit (abcam) for H3K36me3 and one anti-mouse (Santa Cruz) for GAPDH. Images were captured on an ImageQuantTMLAS 4000 (GE Healthcare) for chemiluminescence.

### scRNA-seq data collection and processing

C2C12 (WT and KO) cells were harvested at 80% confluence and washed according to 10X Genomics sample preparation guidelines. Following 10X Genomics Chromium Single Cell 3’ protocol, cells underwent droplet encapsulation and library preparation using Chromium Next GEM Single Cell 3’ GEM Library & Gel Bead Kit v3.1, Chromium Next GEM Chip G, and Dual Index Kit TT Set A. Samples underwent 12 cycles of cDNA amplification and 11 cycles indexing PCRs. Libraries included 1% PhiX spike-in and one sample for WT and KO. They were sequenced on a 150-cycle High v2.5 flow cell loaded on the Illumina NextSeq 500 system. The data were deposited at Gene Expression Omnibus (GEO), accession number GSE304601.

Transcriptomes from approximately 3807 WT and 4982 KO (SETD2-/-) C2C12 cells were processed using the 10x Genomics scRNA-seq pipeline, Cellranger v9.0.1 [90]. Average read depths of 19789 for WT and 13587 for KO were achieved. The Gencode GRCm38 (mm10) primary assembly FASTA and GRCm38 (mm10) primary assembly GTF filtered for protein coding genes were input to ‘cellranger mkref’ to build the reference genome. ‘Cellranger count’ was then used with default arguments to generate gene-barcode matrices. Post-processing was carried out with Seurat, removing genes expressed in fewer than three cells and removing cells expressing fewer than 200 genes [91]. Cells with more than 7% mitochondrial content were removed. WT and KO datasets were normalised using Seurat SCTransform for all analyses besides when raw UMI counts were used to plot overall expression (Fig. 1E) or infer transcriptional bursting parameters.

### Public Histone Modification ChIP-seq dataset collection and processing

We downloaded publicly available ChIP-seq FASTQ for 13 histone modifications and Pol II in murine C2C12 myoblast cells including data from the Encode project, Barbara Wold laboratory (Table S1) [59, 92–95]. For all ChIP-seq data, FASTQ were downloaded from the SRA database. They were all assessed with FASTQC and trimmed using Trimmomatic (arguments: ILLUMINACLIP:Truseq3.fa:2:30:10 LEADING:3 TRAILING:3 SLIDINGWINDOW:4:15) [96]. The minimum length argument was adjusted according to the FASTQ file. Paired-end and single-end FASTQ libraries for each factor were aligned with default settings using Bowtie2 and the Gencode GRCm38 primary genome assembly as reference [97]. Duplicate reads were marked using Picard MarkDuplicates [98]. For the Pol II dataset, the MACS3 callpeak tool was used with arguments (-g mm-q 0.05) to call peaks against its respective input library.

Metagene enrichment profiles were calculated for each histone modification and plotted using Rstudio. We used the Deeptools BamCompare tool (arguments: --effectiveGenomeSize 2652783500 –ignoreDuplicates) to calculate the log2-foldchange of each histone modification against its respective input library [99]. Then ComputeMatrix (arguments: -- beforeRegionStartLength 2000 --afterRegionStartLength 2000 --regionBodyLength 5000 –binSize 10) was used to calculate an average enrichment profile across all non-overlapping protein-coding genes longer than 1.5kb.

### Identifying and Mapping Chromatin States

Chromatin states were modelled with a ChromHMM (v1.25) pipeline using the mm10 assembly [59]. Binaries were generated using default settings and the ChromHMM command, BinarizeBAM, for each sample and their respective input libraries. Models were generated incrementing the number of states modelled by two, from two to twenty states. The models were then analysed to select the optimal number of states.

### State Optimisation

There is no defined method for selecting the optimal number of states to model. The aim of the following analyses is to strike an optimal balance between capturing biologically relevant chromatin states and overfitting the model. Here, we analysed Akaike Information Criterion (AIC) and Bayesian Information Criterion (BIC) to interpret the depreciation of information return when state number increases (Figure S5A). L refers to the maximised likelihood function of the model and K refers to the number of estimated parameters.

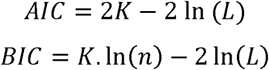

The number of parameters (K) is not explicitly output by ChromHMM, so here we estimated them using the following equation where n is the number of ChIP-seq datasets

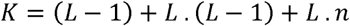

We used ChromHMM’s CompareModels function to compare each state from every model to the most similar state in the twenty-state model. The maximum Pearson’s correlation coefficients for each model were plotted to visualise the redundancy of modelling additional chromatin states (Figure S5B).

To pick an appropriate number of states to model we considered our biological interpretation of each chromatin state, regional enrichment of states, and the previously described analyses.

### Chromatin State Enrichment at Splice Sites

Using junctions extract from the Regtools package, we use the-s XS flag and our WT scRNA-seq BAM dataset as input [62]. From the output we filtered out junctions supported by less than three reads. For each chromatin state, Bedtools intersect was used to count the intersection between consecutive 200 bp bins spanning the genome with active splice sites, the chromatin state, and both features together. Counts were then input into Phyper to assess whether the overlap between active splice sites and each chromatin state is more or less than expected by chance. Bonferroni correction was used for multiple tests.

### Gene Classification Methods

Genes were linked to chromatin states using two methods: (i) by the percentage of their gene body covered by each chromatin state, following separation into bins: 0%; greater than 0% and up to 20%; greater than 20% and up to 40%; greater than 40% and up to 60%; greater than 60% and up to 80%; greater than 80% and up to 100%; and (ii) by assigning the chromatin states at their transcription end sites.

### Expression Analysis

The WT and KO scRNA-seq datasets were used to compare genome-wide gene expression upon abolishing SETD2. We used the three gene classification methods described previously. Gene expression was calculated using Seurat’s SCTransformed scRNA-seq datasets. Mean UMI counts for each gene were plotted to compare expression in WT and KO genotypes. A Mann Whitney U test was used to assess whether differences in gene expression distributions between genotypes were due to chance. Bonferroni correction was used to account for multiple testing.

*Inferring Bursting Parameters*We filtered raw scRNA-seq gene-count matrices for WT and KO by removing genes with zero reads across all cells. We used a previously published modelling strategy [9] to estimate mean expression and burst frequency by means of Markov chain Monte Carlo (MCMC) sampling and calculate burst size and squared coefficient of variation (CV^2^) from samples according to:

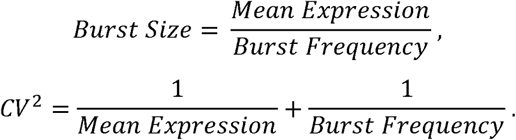

Differences in bursting parameters between conditions (WT vs KO) were assessed using simple linear regression applied to log-transformed parameters, with WT/KO encoded as binary covariate and adjusting for gene-specific log mean expression.

To assess if differences in bursting parameter distributions between genotypes were due to chance for each gene class, Mann Whitney U test was used. Bonferroni correction was used to account for multiple test. Genes were classified by chromatin state as previously described.

### Alternate splicing rMATs

To quantify differential splicing decisions, we used rMATS-turbo (v4.1.2) [101] on pseudo-bulked BAM files generated from scRNA-seq datasets. We note that since single replicates were used, statistical analyses were not possible. We applied the same alignment and post-QC steps using Seurat as described above. We used this tool to quantify the difference in the proportion of reads exhibiting skipped-exon, alternative 5’ splice sites, alternative 3’ splice sites, mutually exclusive exons, or retained introns in absence of SETD2. Due to the 10X Genomics 3’ library kit, we configured rMATs with the following parameters: ‘-t single --readLength 90 --libType fr-unstranded --allow-clipping’.

Next, we used the mutually exclusive exon (MXE) pairs identified by rMATs to investigate the effect of H3K36me3 enrichment in splice site decision making. First, we calculated the RNA read density using our pseudo-bulked scRNA-seq wild-type bam file, normalising for exon length. Read counts were calculated by converting our MXE annotations to SAF for use with subread featurecounts. To assess H3K36me3 enrichment over input, deeptools ‘bamCompare’ was used with the parameters ‘-binSize 10 –operation log2 –scaleFactorsMethod readCount –pseudoCount 1 –ignoreReplicates’ [102]. A blacklist was applied to remove regions with low mappability. Next, we passed the output from bamCompare to ‘multiBigWigSummary BED-file’ to assess average H3K36me3 enrichment for every MXE exon in our annotation file. Aware of the bias of using 3’ scRNA-seq data, we sought to mitigate this by removing MXE pairs where the RNA enrichment was greater in downstream exons when comparing MXE pairs. We also only considered MXE pairs with enrichment scores greater than 0 for both ChIP-seq and RNA-seq data. Spearman Rank correlation test was used to assess whether H3K36me3 enrichment between MXE pairs influences RNA enrichment and thus splice site decision making.

### Velocyto and evaluating splicing efficiency

To evaluate global pre-mRNA splicing efficiency, we generated spliced and unspliced counts using the ‘Velocyto run’ command-line tool. Velocyto generates a data object with sequencing read count data in 3 layers: spliced (reads mapping to only exonic regions), unspliced (reads containing intronic regions), and ambiguous (reads that could not be assigned) [103]. Since we present splicing efficiency as a ratio, normalisation for library depth was unnecessary. Global splicing efficiency was calculated per cell, based on the ratio of total unspliced reads to spliced reads. The differences were then visualised using a violin plot with Mann-Whitney U test.

## SUPPLEMENTARY INFORMATION

**Table S1.** Publicly sourced C2C12 myoblast ChIP-seq datasets. Including histone modifications, RNA polymerase II, and their respective input libraries.

| ChIP-seq | Accession ID | Sample ID |
| --- | --- | --- |
| H3ac (ab 06-599) | GSE36023 | GSM918422 |
| H3K18Ac | GSE25308 | GSM721302 |
| H3K27ac | GSE161054 | GSM4889016 |
| H3K27me3 | GSE36023 | GSM918408 |
| H3K36me3 | GSE36023 | GSM918417 |
| H3K4me1 | GSE25308 | GSM721288 |
| H3K4me2 | GSE25308 | GSM721290 |
| H3K4me3 | GSE25308 | GSM721292 |
| H3K79me2 | GSE36023 | GSM918411 |
| H3K79me3 | GSE36023 | GSM918410 |
| H3K9Ac | GSE25308 | GSM721300 |
| H3K9me3 | GSE65493 | GSM1598791 |
| H4K12Ac | GSE25308 | GSM721304 |
| Input Control | GSE65493 | GSM1716540 |
| Input Control | GSE161054 | GSM4889019 |
| Input Control 32bp | GSE36023 | GSM918419 |
| Input Control 50bp | GSE36023 | GSM918421 |
| MNAse digested<br>input | GSE25308 | GSM721308 |
| RNA Polymerase II | GSE25308 | GSM721286 |
| Sonicated_input_MB | GSE25308 | GSM721306 |

---

**Figure S1.**
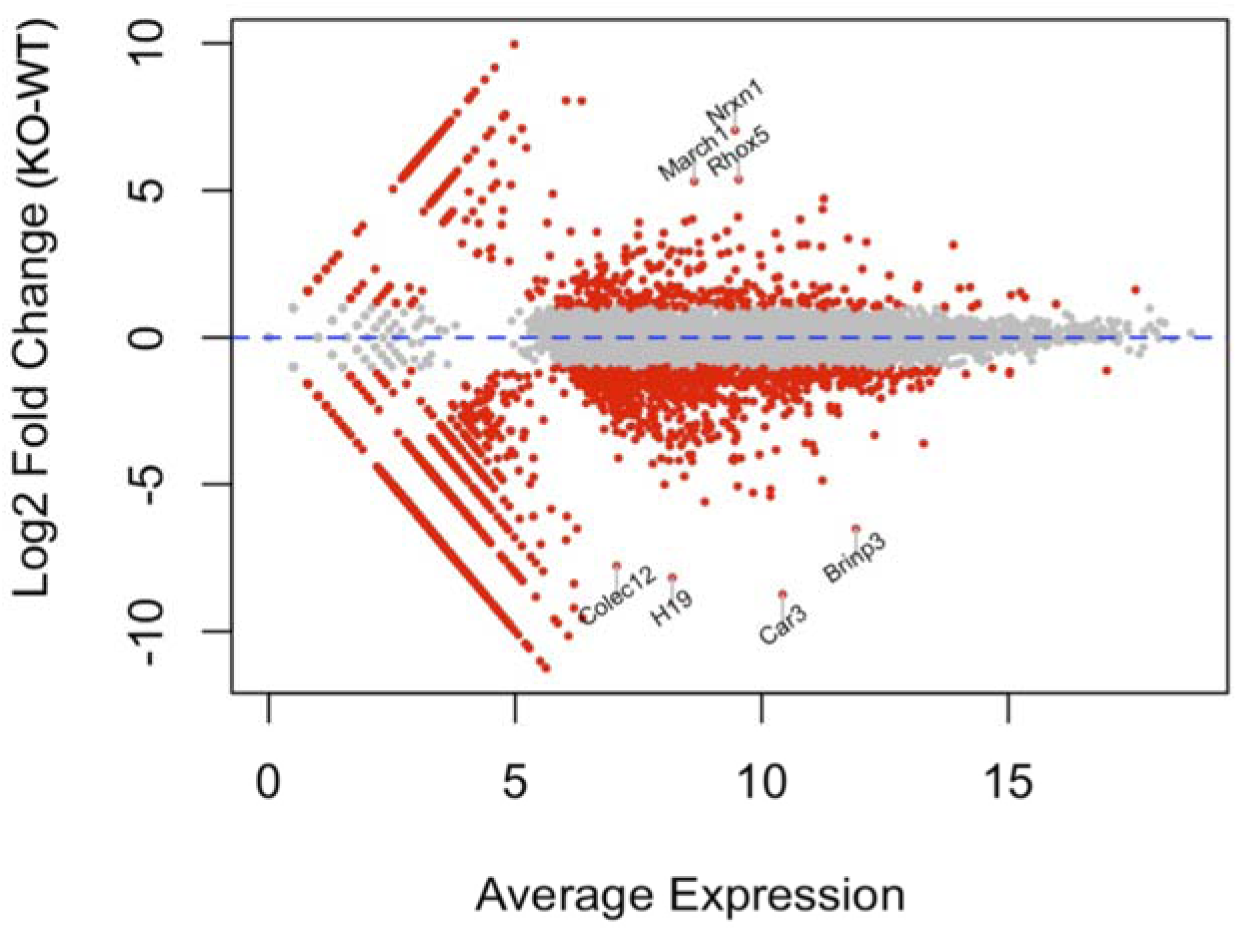
MA plot of gene expression in KO vs WT cells.

**Figure S2.**
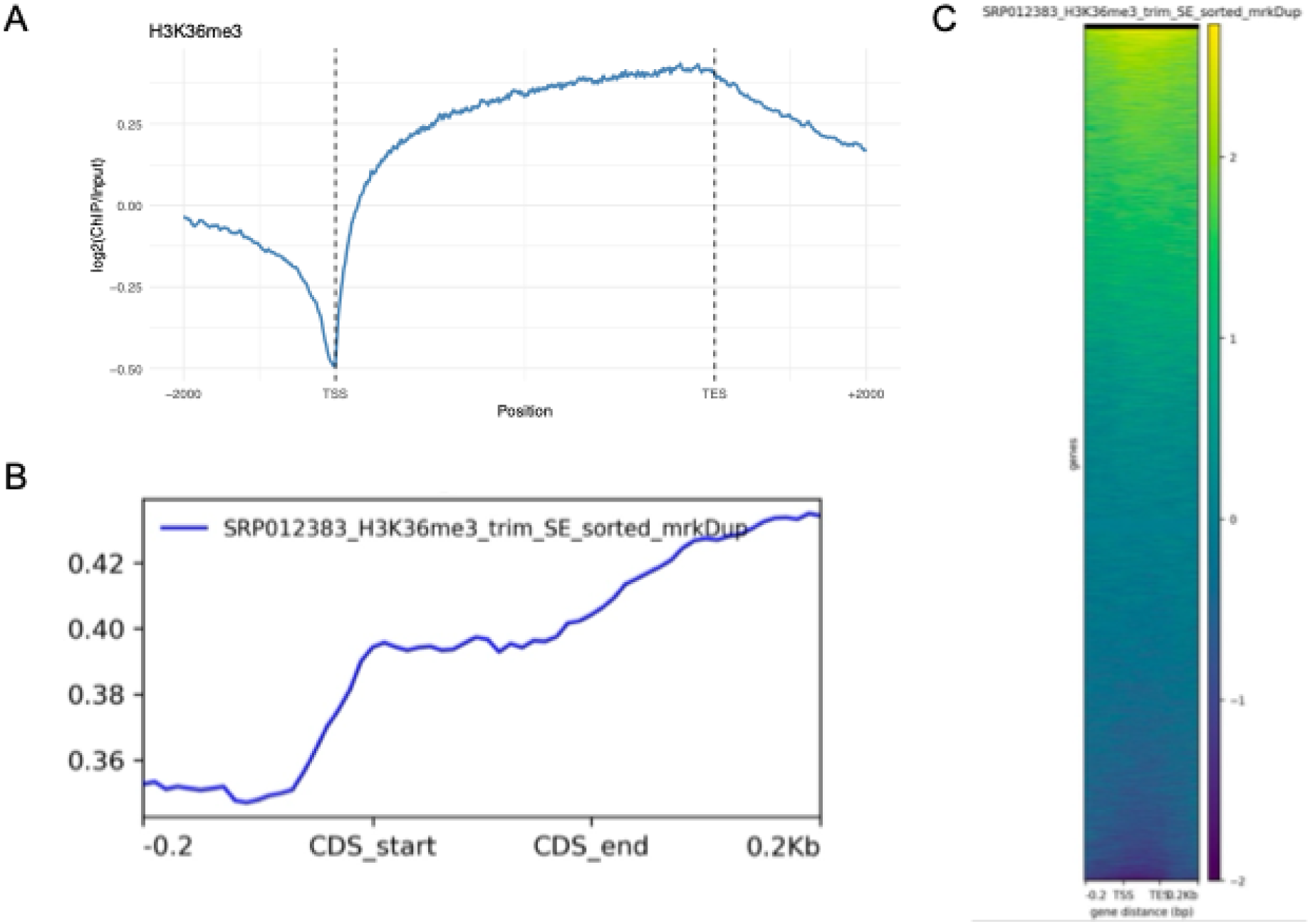
H3K36me3 mark distribution in C2C12 cells. **A**, Metagene profile at gene body. **B**, Metagene profile at CDS. **C**, as B, shown as heatmap that resolves individual transcripts. Dataset GSE36023.

**Figure S3.**
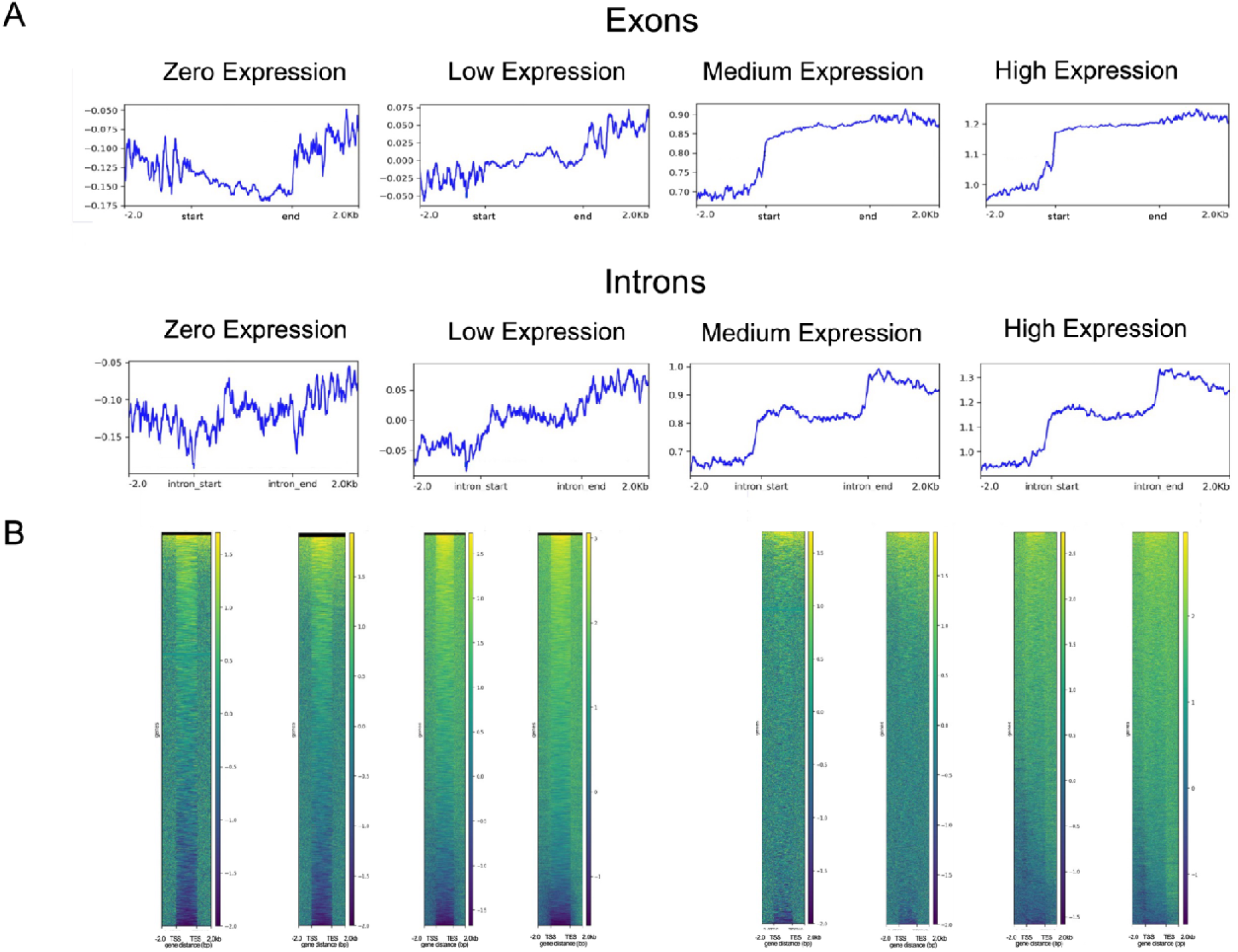
Intragenic H3K36me3 distributions stratified by gene expression level. A, Average densities of H3K36me3 marks in the CDS (top row) and introns (bottom row) relative to inputs, for genes classified into four different expression level categories as indicated. **B**, as A, shown as heatmaps that resolve individual transcripts (CDS, left; introns, right). Dataset GSE36023.

**Figure S4.**
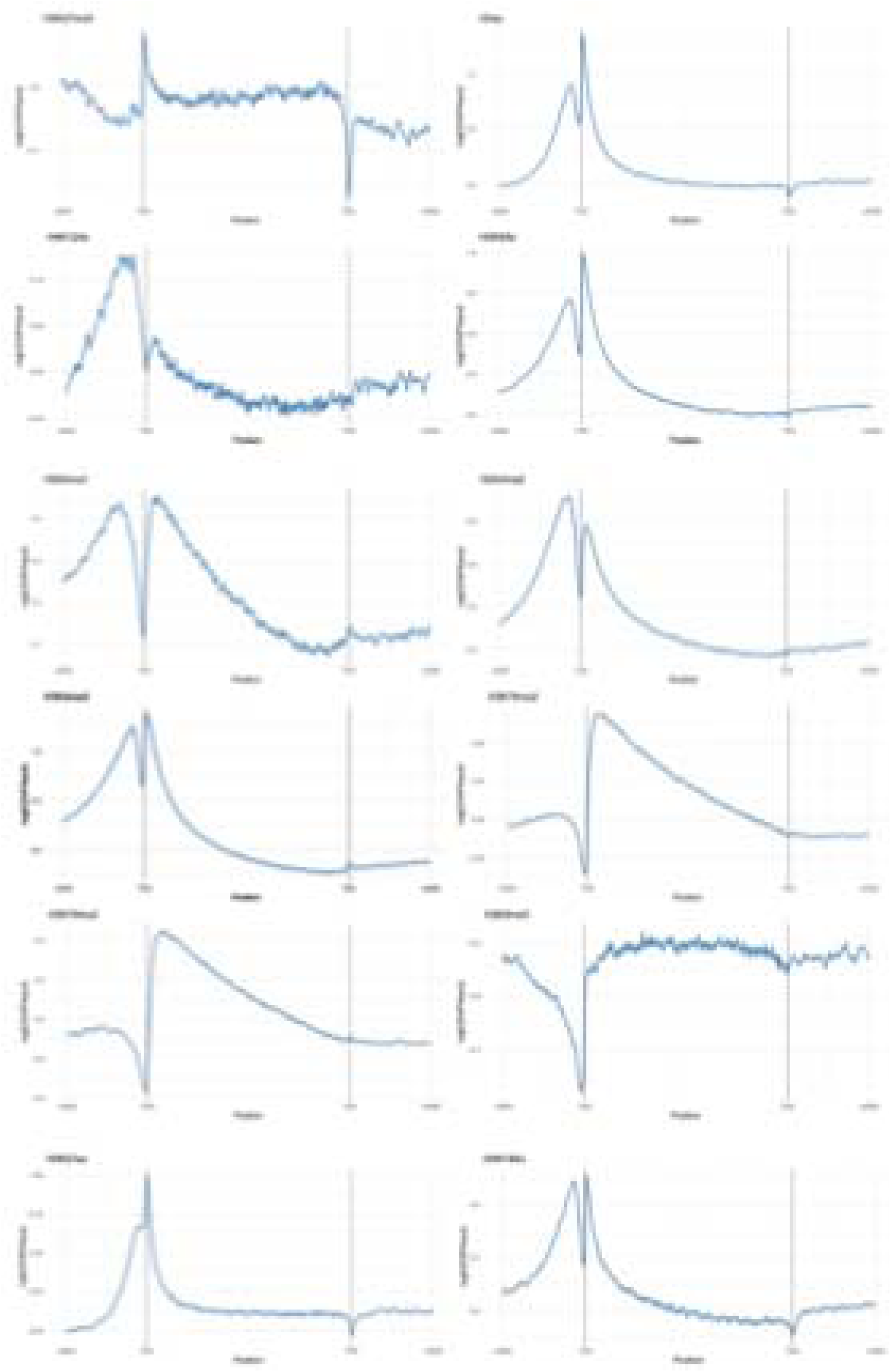
Metagene profiles for histone modifications.

**Figure S5.**
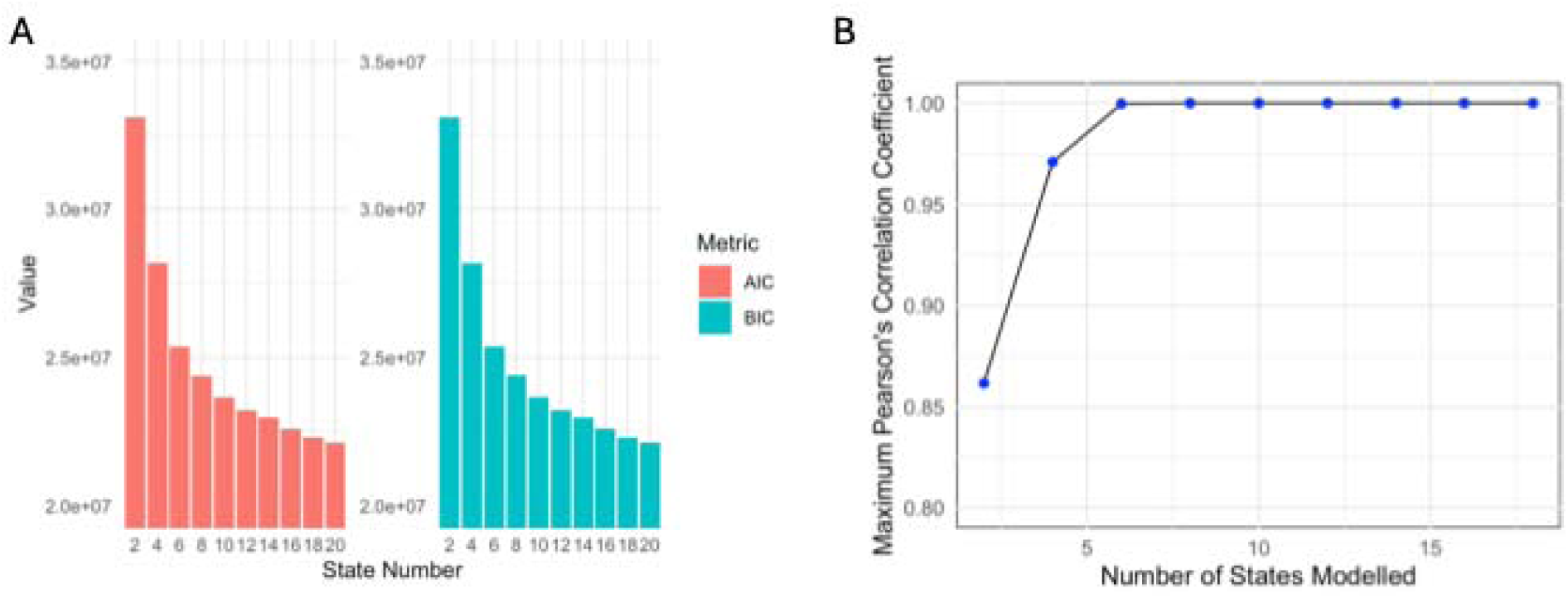
Chromatin state number metrics. **A**, Akaike Information Criterion (AIC) and Bayesian Information Criterion [15] as functions of the number of states. **B**, Assessment of redundancies among different numbers of ChromHMM states. Pairwise Pearson correlation coefficients were calculated for all states of a 20-state reference model and smaller ones and the maximum values in each case plotted as shown.

**Figure S6.**
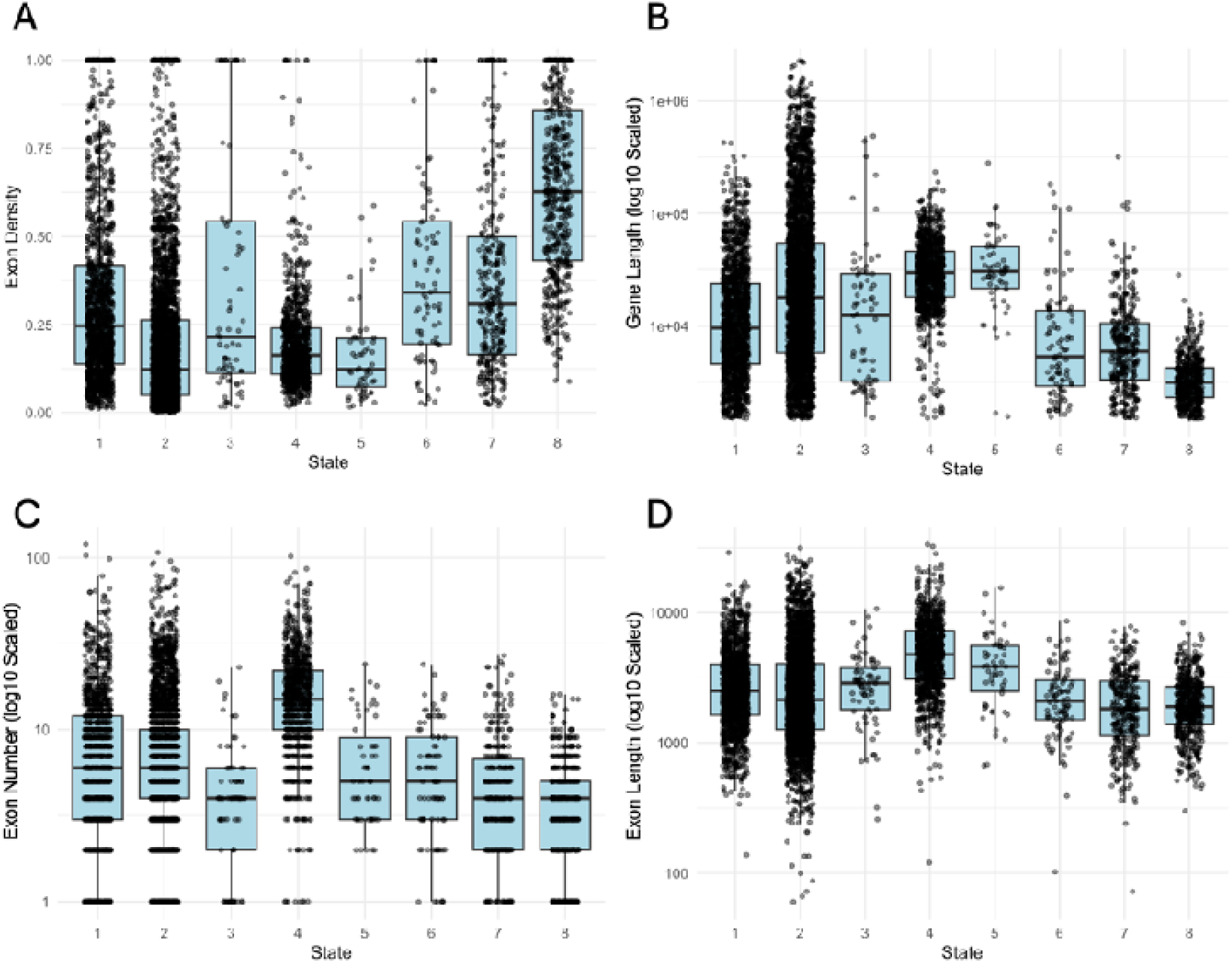
Genomic features of high coverage genes for each chromatin state. Genes were classified under each chromatin state if they exhibited greater than 80% coverage for the given state. Features for all genes were calculated from Mus musculus (mm10) annotations. Exon density was calculated from exon length (bp) divided from gene length (bp). Sample size for each state; state 1 (n = 1297), state 2 (n = 3144), state 3 (n = 67), state 4 (n = 685), state 5 (n = 41), state 6 (n = 73), state 7 (n = 259), and state 8 (454).

**Figure S7.**
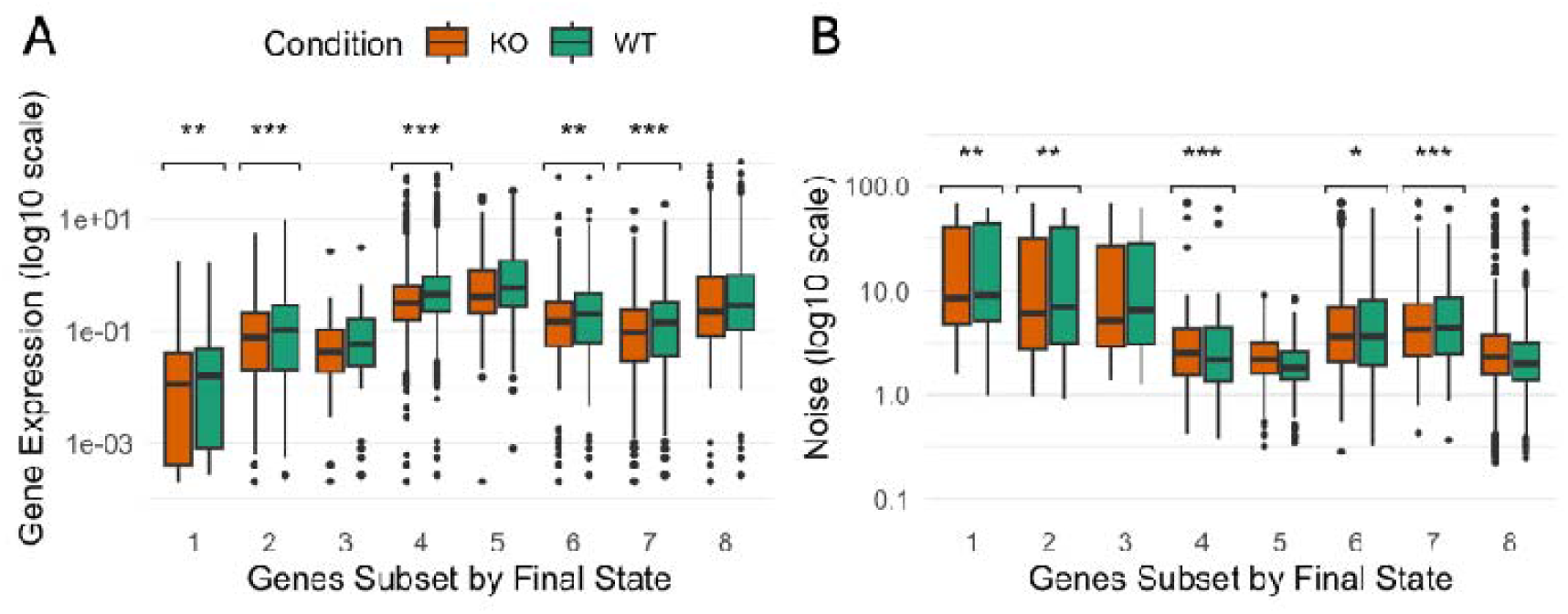
The effect of H3K36me3 loss on gene expression and noise, stratified by the final state in each gene. Genes were classified according to their final chromatin state (1 to 8; Fig. 5). **A**, Mean expression levels were calculated based on UMI counts from scRNA-seq data. **B**, as A, for noise (CV^2^). Genotypes were compared using Mann Whitney U test with Bonferroni correction (* < 0.05, ** < 0.01, *** < 0.001). Number of genes sampled for each state (WT, KO): state 1 (n = 436, 333), state 2 (n = 2085, 1785), state 3 (n = 112, 90), state 4 (n = 3503, 3463), state 5 (n = 141, 138), state 6 (n = 880, 810), state 7 (n = 1414, 1287), and state 8 (n = 596, 570).

**Figure S8.**
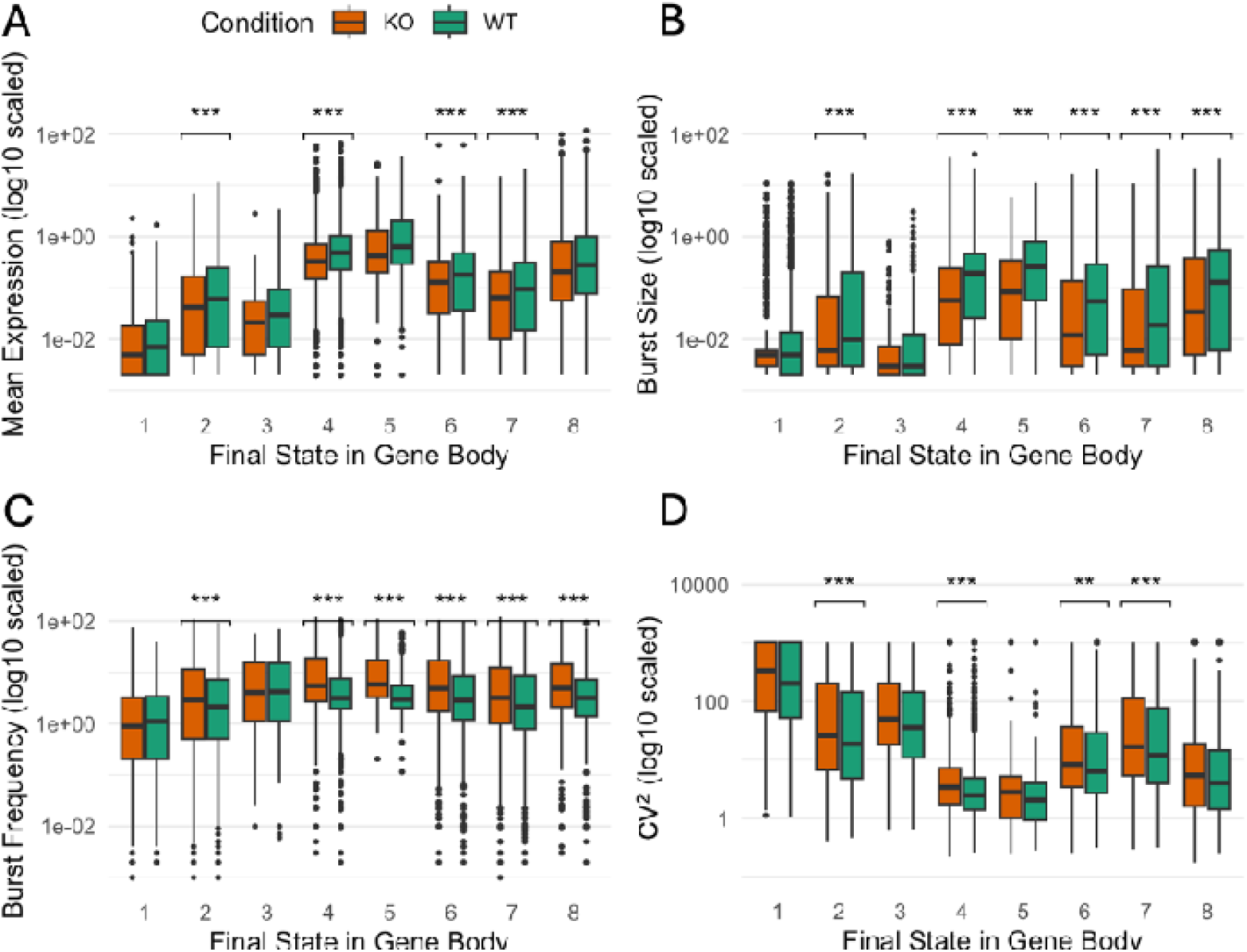
The effect of H3K36me3 loss on bursting parameters, stratifying genes by their most 3’ chromatin state. Mean expression and burst frequency were estimated using a Markov chain Monte Carlo (MCMC) approach on scRNA-seq count data. Plots compare wild-type (WT) and SETD2 knockout (KO) gene regarding **A**, mean expression, **B**, burst size, **C**, burst frequency, and, **D**, cumulative variation (CV^2^). Genotypes were compared using Mann Whitney U test with Bonferroni correction (* < 0.05, ** < 0.01, *** < 0.001). Number of genes sampled for each state (WT, KO): state 1 (n = 728, 561), state 2 (n = 2616, 2337), state 3 (n = 166, 148), state 4 (n = 3721, 3692), state 5 (n = 149, 147), state 6 (n = 1001, 947), state 7 (n = 1729, 1587), and state 8 (n = 681, 657). Burst size = mean expression / burst frequency. CV^2^ = (1/mean expression) + (1/burst frequency).

